# NAD homeostasis maintained by NMNAT2 supports vesicular glycolysis and fuels fast axonal transport in distal axons of cortical glutamatergic neurons in mice

**DOI:** 10.1101/2022.02.06.479307

**Authors:** Sen Yang, Zhen-Xian Niou, Andrea Enriquez, Jacob LaMar, Jui-Yen Huang, Karen Ling, Paymaan Jafar-Nejad, Jonathan Gilley, Michael P. Coleman, Jason M. Tennessen, Vidhya Rangaraju, Hui-Chen Lu

**Affiliations:** The Linda and Jack Gill Center for Biomolecular Sciences, Indiana University, Bloomington, IN 47405; Department of Psychological and Brain Sciences, Indiana University, Bloomington, IN 47405.; Program in Neuroscience, Indiana University, Bloomington, IN 47405; Max Planck Florida Institute for Neuroscience, Jupiter, FL, USA; Neuroscience Drug Discovery, Ionis Pharmaceuticals, Inc., 2855 Gazelle Court, Carlsbad, CA 92010, USA; Department of Clinical Neuroscience, Cambridge University, Cambridge, United Kingdom; Department of Biology, Indiana University, Bloomington, IN 47405, USA

## Abstract

**Background:** Bioenergetic maladaptations and axonopathy are often found in the early stages of neurodegeneration. Nicotinamide adenine dinucleotide (NAD), an essential cofactor for energy metabolism, is mainly synthesized by Nicotinamide mononucleotide adenylyl transferase 2 (NMNAT2) in CNS neurons. NMNAT2 mRNA levels are reduced in the brains of Alzheimer’s, Parkinson’s and Huntington’s disease. Here we addressed whether NMNAT2 is required for axonal health of cortical glutamatergic neurons, whose far-projecting axons are vulnerable to neurodegenerative conditions. We also tested if NMNAT2 maintains axonal health by ensuring proper axonal ATP levels for axonal transport, a critical function of axons.

**Methods:** We generated mouse and cultured neuron models to determine the impact of NMNAT2 loss from cortical glutamatergic neurons on axonal transport, energetic metabolism, and morphological integrity. In addition, we determined if exogenous NAD supplementation or inhibiting NAD hydrolase sterile alpha and TIR motif-containing protein 1 (SARM1) prevented axonal deficits caused by NMNAT2 loss. Our study used a combination of genetic, molecular biology, immunohistochemistry, biochemistry, fluorescent time-lapse imaging, live imaging with optical sensors, and anti-sense oligos application.

**Results:** We provide *in vivo* evidence that NMNAT2 in cortical glutamatergic neurons is required for axonal survival. Using *in vivo* and *in vitro* studies we demonstrate that NMNAT2 protects axons by ensuring the proper NAD-redox potential in distal axons of cortical neurons to support glycolysis on vesicular cargos, thus ensuring “onboard” ATP production fueling axonal transport. Exogenous NAD^+^ supplementation to NMNAT2 KO cortical neurons restores glycolysis and resumes fast axonal transport. Finally, we demonstrate both *in vitro* and *in vivo* that reducing the activity of SARM1, an NAD degradation enzyme, can reduce axonal transport deficits and suppress axon degeneration in NMNAT2 KO neurons.

**Conclusion:** NMNAT2 ensures axonal health by maintaining NAD redox potential in distal axons to ensure efficient vesicular glycolysis required for fast axonal transport.

## Background

Glucose is the primary energy source in the brain [1, 2]. Recent human studies highlight bioenergetic mal-adaptations in the brain during neurodegenerative disorders of aging (NDA) [3], such as Alzheimer’s disease (AD). In AD, the white matter of the brain [4], which comprises long-range axonal tracts, exhibits glucose hypometabolism. Axons, the longest and the most complex subcellular compartments of neurons [5–7], are particularly vulnerable to neurodegenerative conditions [8–10]. Key questions concerning glucose metabolism in axons during NDAs include: Does glucose hypometabolism in long-range axons cause axonopathy, an early sign of neurodegeneration [11, 12]? And what are the key steps in maintaining axonal energetics to ensure axonal function and health?

Nicotinamide adenine dinucleotide (NAD) is a cofactor essential for energy metabolism. The ratio of oxidized (NAD^+^) to reduced (NADH) form of NAD, called the NAD redox potential, is a pivotal driver of glycolysis, tricarboxylic acid (TCA) cycle, and oxidative phosphorylation (OXPHO) [13]. Nicotinamide mononucleotide adenylyl transferase 2 (NMNAT2) is the major NAD synthesizing enzyme in CNS neurons [14–16] and has been shown to be a key axonal maintenance factor [17–20]. mRNA levels of NMNAT2 are significantly reduced in the brains of AD, Parkinson’s disease (PD), and Huntington’s disease (HD) patients [21–23]. Notably, upregulating NMNAT2 and other NMNATs is protective in multiple neurodegenerative disease models [23, 24]. Whether NMNAT2 maintains axonal health by contributing to axonal bioenergetics has not been examined.

Previous studies showed that NMNAT2 is localized to Golgi-derived vesicles in axons and undergoes fast, bi-directional axonal transport in superior cervical ganglia neurons [25, 26]. Axonal transport between the cell body and axonal terminals is critical for neurotransmission, synaptic plasticity [27], and neuronal health [28]. To achieve fast and continuous movement of cargo along highly arborized axons across long distances [29–31], the bioenergetic machinery must be locally mobilized to provide ATP. It has been demonstrated that glycolytic enzymes specifically attach to fast moving vesicular cargos to fuel fast axonal transport [32, 33]. Surprisingly, ultra-resolution electron microscopy studies find that mitochondria in axons are often small in size and sparsely distributed [34, 35]. Moreover, *in vitro* evidence suggests that glycolysis, rather than mitochondrial OXPHO, supplies constant energy to fuel fast axonal transport [32]. Moving one vesicular cargo at physiological speed consumes ∼200 molecules of ATP molecules per second [36], while each cycle of glycolysis only yields 2 molecules of ATPs. Thus, vesicular glycolysis in axons has to turnover rapidly to support fast axonal transport. In this study, we aimed to test the hypothesis that NMNAT2 maintains the NAD redox potential necessary to drive glycolytic ATP synthesis for fast axonal transport.

We employed both animal (*in vivo*) and culture (*in vitro*) model systems to determine: (1) the requirement of NMNAT2 for the health of long-range axons of glutamatergic neurons; (2) whether NMNAT2 is essential for fast axonal transport; (3) whether NMNAT2 ensures the proper NAD^+^/NADH redox potential for glycolysis on vesicular cargos; and (4) the efficacy of NAD supplementation in reversing the axonal phenotypes caused by NMNAT2 loss. Our results reveal the importance of NMNAT2 to maintain the axonal NAD redox potential and drive the onboard glycolysis necessary for fast axonal transport; we also demonstrate that NMNAT2 reduction increases the vulnerability of distal axons.

## Methods

### Mice

The generation of NMNAT2^f/f^ mice carrying a Cre recombinase-dependent gene trap cassette within intron 1 of NMNAT2 gene has been described [37]. Here we generated NMNAT2 conditional knockout (cKO) mice by crossing NMNAT2^f/f^ mice with NEX-Cre mice [38] to delete NMNAT2 in cortical glutamatergic neurons after embryonic day 11.5 (E11.5). NMNAT2 cKO mice were smaller in size and exhibited an ataxia phenotype with wobbling movements. Thus, 5 grams of DietGel 76A (Clear H_2_O, Westbrook, ME, USA) were given daily after animals were weaned. NEX-Cre positive NMNAT2^f/+^ and NMNAT2^+/+^ and NEX-Cre negative NMNAT2^f/f^ littermates were used as controls for NEX-Cre positive NMNAT2^f/f^. The generation and genotyping of NMNAT2-Blad mice have been described previously [39]. In these mice, the NMNAT2 null mutation was generated by transposon-mediated gene-trap mutagenesis. Sarm1 KO (S^null^/S^null^) mice [40] were crossed with NMNAT2 cKO mice to generate cKO;S^null^/+ and cKO; S^null^/S^null^ transgenic mice. Littermate controls include NEX-Cre/+;NMNAT2^f/+^;S^null^/+, NMNAT2^f/+^;S^null^/+; NEX-Cre/+;NMNAT2^f/+^;S^null^/ S^null^, and NMNAT2^f/f^;S^null^/ S^null^. Both male and female mice were used for all experiments. All mice were housed under standard conditions with food and water provided ad libitum and maintained on a 12 h dark/light cycle. Mice were housed and used in compliance with the NIH Guidelines for the Care and Use of Laboratory Animals and were approved by the institutional animal care and use committees at Indiana University and Max Planck Florida Institute for Neuroscience.

### Genotyping

For mice, ear lysates were prepared by immersing the tissue derived from a small ear clip in digestion buffer (50 mM KCl, 10 mM Tris–HCl, 0.1% Triton X-100, 0.1 mg/ml proteinase K, pH 9.0), vortexing gently, and then incubating for 3 h at 60°C to lyse the cells. For embryos, the same digestion buffer with 0.2 mg/ml proteinase K was used, and embryonic tail lysates were incubated at 60°C for 15 min with 1500 rpm shaking in a thermoshaker. These lysates were then heated to 95°C for 10 min to denature the proteinase K (Thermo Scientific, Rockford, IL, USA), and centrifuged at 16,100 g for 15 min. The supernatants were used as DNA templates for polymerase chain reactions (PCRs, EconoTaq Plus Green 2X mater mix, Lucigen, Middleton, WI, USA). For embryonic genotyping during primary culture preparation, a QIAGEN Fast Cycling PCR Kit was used. The sequences of the primers used for genotyping are listed in the reagent table. For NMNAT2^f/f^ line: primer A and B detect WT allele; primer B and C detect NMNAT^f^ allele. For NEX-Cre line: primer Cre484 and Cre834 detect Cre positive allele. For NMNAT2-Blad line: primer A and B detect WT allele; primer R3 and Rf detect KO allele. For SARM1 KO line: primer WT-R and Sarm1-common detect WT allele, primer Sarm1-common and Mut-R detect KO allele.

### Immunohistochemistry for brain sections

Mice were anesthetized and perfused with 4% paraformaldehyde (PFA) in PBS. Brains were harvested, post-fixed with 4% PFA in PBS overnight at 4°C, and then rinsed with PBS. Free floating brain sections were then prepared by sectioning these fixed brains into 40-μm-thick sections in the coronal plane with a Leica VT-1000 Vibrating microtome or Sliding Microtome SM-2000R (Leica Microsystems).

For immunohistochemistry, sections were permeabilized with 0.3% Triton X-100 in PBS for 20 min at room temperature, incubated with blocking solution (3% normal goat serum prepared in PBS with 0.3% Triton X-100) for 1 h, and then incubated with primary antibodies diluted in blocking solution overnight at 4°C. The next day, sections were washed with 0.3% Triton PBS 3 times and then incubated with the secondary antibodies diluted in blocking solution at 4°C for 2 h. After the incubation, samples were washed with 0.3% Triton PBS 3 times. Draq5 (1:10,000 dilution, Cell Signaling) or 4°,6-diamidino-2-phenylindole (DAPI, 5 μg/ml, Invitrogen) were added during the first wash step to visualize nuclei. Dako mounting medium was used to mount the brain sections.

### Microscopy, imaging, and data analysis for brain sections

Bright-field images were obtained with Zeiss SteREO Discovery.V8 Microscope (Carl Zeiss Microscopy). Confocal fluorescent images were taken by a Leica TCS SPE confocal microscope (Leica DM 2500) with a 10x objective lens (0.3 N.A.) or a 40x (0.75 N.A.) objective lens. Some confocal images were taken with a Nikon A1R laser scanning confocal (Nikon A1) with a 10x (0.5 N.A.), or a 60x (1.4 N.A.) oil objective lens. DAPI/Draq5 immunofluorescence was used to identify comparable anatomical regions across different brain sections. A minimum of three sections were imaged per mouse, and each anatomical region was imaged with both side of the cortices and comparable across all animals. Images for a particular staining were acquired with identical imaging parameters selected to minimize signal saturation in all experimental groups. The thickness of the corpus callosum was measured from the dorsal to ventral edges of NFM positive axonal tracts in coronal plane brain sections, while the thickness of the primary somatosensory cortex was measured from the pial surface to white matter regions using image J “Straight” function. The total pixel values of APP signals and area sizes were measured by Image J to acquire APP signal densities.

### NMNAT2 in situ hybridization

In situ hybridization on embryonic sections was done by the Baylor College of Medicine Advanced Technology Core Labs. The experiment was performed using a high-throughput automated platform. Cryostat sections of E14.5 embryos were placed on a standard microscope slide that was subsequently incorporated into a flow-through chamber. The chamber was then placed into a temperature-controlled rack and the required solutions for pre-hybridization, hybridization, and signal detection reactions were added to the flow-through chamber with an automated solvent delivery system. Details can be found in a previous publication [5]. The RNA probe was generated using the sequence from the Allen Brain Atlas database. Images were acquired by Leica CCD camera with a motorized stage. Multiple images were collected from the same section. Individual images were stitched together to produce a mosaic representing the entire section.

### Primary cortical neuronal cultures

To culture NMNAT2 KO neurons, E15.5-16.5 embryos were harvested from NMNAT2-Blad heterozygous (HET) female mice mated to NMNAT2-Blad HET male mice. Embryos of both sexes were included for culture preparations. The cortical tissue of each embryo was dissected and then dissociated individually using the Worthington Papain Dissociation Kit (Worthington Inc.) according to the manufacturer’s protocol. Genotyping was performed before the completion of papain dissociation. Cortices from multiple NMNAT2-Blad wildtype (WT) and HET embryos were combined as control group for dissociation and plating; the cortices from multiple NMNAT2-Blad homozygous embryos were pooled together as KO group for dissociation and plating. NMNAT2^f/f^ neuron cultures were prepared from cortical tissue dissected from E15.5-16.5 NMNAT2^f/f^ embryos harvested from NMNAT2^f/f^ female mice mated with NMNAT2^f/f^ male mice. After plating, cortical neurons were maintained in Neurobasal Media (Gibco™, ThermoFisher Scientific) supplemented with 2% B-27 (Gibco™), 2 mM GlutaMAX™ (Gibco™) and 100 U/ml penicillin-streptomycin (Gibco™), incubated at 37°C with 5% CO2 and appropriate humidity. One third of the media was replenished every 3 days.

### Immunocytochemistry and confocal imaging

About 2.5*10^5^ cortical neurons were plated on 12 mm diameter coverslips coated with poly-D-lysine (PDL) in 24-well plates (1.3*10^3^ cells/mm^2^). At indicated days *in vitro*, cortical neurons were first fixed by 4% PFA and 4% sucrose in PBS for 20 min, incubated 60 min in blocking buffer (0.1% Triton X-100 and 5% Goat Serum in PBS), and then incubated with primary antibody containing blocking buffer overnight at 4°C on a gentle rotating platform. Samples were then washed with 0.1% Triton in PBS 3 times and incubated with secondary antibody containing blocking buffer for 2 h. After the incubation, samples were washed with 0.1% Triton in PBS 3 times. Coverslips were then taken out to mount with ProLong™ Gold antifade mounting medium with DAPI.

Fluorescent confocal images were taken by a Leica TCS SPE confocal microscope using a 63x (1.4 N.A.) oil objective lens or by a Nikon A1R-HD25 laser scanning confocal microscope with an Apo Lambda S 60x (1.4 N.A.) oil objective lens. Z-stack images were taken with 0.5 μm Z-step size to cover the whole depth of the cultured neurons. Images were acquired with identical imaging parameters chosen to optimize signal to noise ratio and avoid saturated pixels in any experimental groups. To quantify APP or TUJ1, 3-5 regions were randomly selected per coverslip for imaging. Each image accounts for one data point in quantification. Two coverslips per group were imaged for each batch of culture. To sample a larger area for imaging MAP2 and βIII-tubulin staining for gross neurite area quantification, the “large image stitching” function in Nikon NIS-Elements software with 5-10% overlap was used to tile 4 adjacent fields of view together as one image.

### Quantification of APP accumulation and gross neurite area in neuronal culture

ImageJ was used for quantification of the maximum intensity projected images. APP signal in somata and dendrites was manually excluded, then the default threshold method was applied to select the top 10% tail of total pixels, and the “analyze particles” function was applied to select particles of area larger than 0.9 μm^2^ which were identified as “accumulated APP”. The total area of “accumulated APP” was calculated in each image to evaluate the phenotype. MAP2 positive areas were identified by means of an auto-thresholding method and measured for each image. A MAP2 positive area was selected by the “create selection” function after thresholding, then restored and deleted from the βIII-tubulin channel. Therefore, the remaining βIII-tubulin signal solely represented axonal regions. Similarly, mean threshold was applied to measure the βIII-tubulin positive area. Live cells were distinguished from dead cells by DAPI-stained nuclei morphology, and the number of live cells was counted for each image. MAP2 and βIII-tubulin areas were normalized by dividing by the live cell number in each image. We observed a skew in the data if the cell density was dramatically different between Ctrl and KO. Therefore, only images with a live cell number between 150 to 300 within 0.31 mm^2^ image area (485-970 cells/mm^2^) were used for the analysis.

### Plasmid DNA constructs

pEGFP-n1-APP was a gift from Zita Balklava and Thomas Wassmer (Addgene plasmid # 69924; http://n2t.net/addgene:69924; RRID: Addgene_69924) [41]. pEGFP-c1-SNAP25, pEGFP-n1-SYPH, and pmCherry-n1-NMNAT2 were gifts from Michael Coleman and Jonathan Gilley [25]. pCX-EGFP was a gift from Matthew Neil Rasband [42]. pLV-mitoDsRed was a gift from Pantelis Tsoulfas (Addgene plasmid # 44386; http://n2t.net/addgene:44386; RRID: Addgene_44386) [43]. pcDNA3.1-SoNar and pcDNA3.1-cpYFP were gifts from Yang Yi [44]. pCMV-MitoVenus was a gift from Lulu Cambronne [45]. pCAG-mCherry was a gift from Ken Mackie. pcDNA3-Syn-ATP and pcDNA3-Cyto-pHluorin were from Vidhya Rangaraju [33].

### Transfection in neuronal culture

For lentiviral transduction in NMNAT2^f/f^ neurons, a lentivirus expressing copGFP or iCre under elongation factor 1 alpha (ELF1alpha) promoter was applied at 2 multiplicity of infection (MOI) at 9 days in vitro (DIV9).

For lipofectamine transfection, about 1.6*10^6^ cortical neurons were plated onto PDL (Poly-D-Lysine)-coated MatTek dish (P35G-1.5-20-C) (1.66*10^3^ cells/mm^2^). For APP-EGFP and SNAP25-EGFP transport imaging, cortical neurons were transfected 16 to 20 h prior to the time-lapse imaging experiments using Lipofectamine 3000 (Thermo) following manufacturer’s instruction. For MitoDsRed transport imaging, cortical neurons were co-transfected with pCX-EGFP and pLV-mitoDsRed 36 to 48 h before live imaging using Lipofectamine 3000. Before adding the DNA-lipofectamine mixture, half of the conditioned culture medium was removed and saved for later. Two hours after incubation with DNA-lipofectamine mixture, one third of the remaining medium was further removed and 2 ml of culture medium (1:1 of conditional medium and fresh medium) was added back immediately. For live imaging, all culture medium was replaced by 2-3 ml of Hibernate E low fluorescence buffer (BrainBits) supplemented with 2mM GlutaMAX™ to maintain the ambient pH environment.

For magnetofection, about 2.56*10^5^ cortical neurons were plated onto the center of a PDL-coated coverglass region of MatTek dish (P35G-1.5-14-C) (1.66*10^3^ cells/mm^2^). For Syn-ATP imaging and Cyto-pHluorin imaging, cortical neurons were transfected 24 h prior imaging using Combimag (OZ biosciences) and Lipofectamine 2000 (Invitrogen) according to the manufacturer’s instruction. Briefly, 0.5 μg DNA was incubated with 0.5 μl lipofectamine in 50 μl transfection medium (Neurobasal media supplemented with 2 mM Glutamax without B27 or antibiotics) for 5 min, then mixed with 0.5 μl Combimag diluted in another 50 μl transfection medium for 10 min. The DNA-lipofectamine-Combimag mixture was further diluted in 125 μl of transfection medium. Then, the conditioned medium from cultured neurons was harvested and the neurons were immediately rinsed twice in warm transfection medium. Transfection medium used for rinsing was removed and 150 μl of DNA-lipofectamine-Combimag mixture was added to the neurons. Neurons were placed on a magnetic plate for 20 min inside a 37°C and 5% CO_2_ incubator. After 20 min, neurons were rinsed once in warm transfection medium and replaced with the previously harvested warm, conditioned medium.

### Axonal transport time-lapse video microscopy and movie analyses

Live imaging of neuronal cultures was carried out using an Olympus OSR Spinning Disk Confocal microscope (CSU-W1, Yokogawa) connected to a Hammamatsu Flash 4 V2 camera. Temperature was maintained at 37°C by a Tokai Hit Stage Top Incubation system. For APP-EGFP and SNAP25-EGFP transport recording, movies were taken in axonal segments at least 400 μm away from the soma (defined as distal axon) and axonal segments within 200 μm of the soma (defined as proximal axon) and captured at a rate of 5 frames/s for a total recording time of 60 s with a 63x oil objective (1.46 N.A.). For MitoDsRed transport imaging, neurons were co-transfected with EGFP, and the distance from axonal segment of interest to soma was determined in the GFP channel under the eyepiece. Mitochondria transport videos were taken in the distal region at a 3 second time interval for a total recording time of 15 min with a 63x oil objective (1.46 N.A.). A z-stack with 1 to 1.5 μm z-step size was taken for each time point to cover all the mitochondria within the imaged axonal segment.

Kymographs were generated using ImageJ (http://rsb.info.nih.gov/ij) with “Velocity_Measurement_Tool” macro (http://dev.mri.cnrs.fr/projects/imagej-macros/wiki/Velocity_Measurement_Tool). Quantification of velocity was manually performed by following the trajectory of each particle with an angle larger than 0° relative to the time axis. Stationary and repetitive bidirectional-moving particles were excluded for velocity measurements. If an anterograde or retrograde moving particle stopped for a short while during the imaging and then continued moving, the paused portion was included in the velocity calculation. Although a portion of the axon could be out of focus, resulting in a discontinuous trajectory, the discontinuous trajectory was still traced as the same if the slope and timing matched. All the traced trajectories were saved in “ROI manager” in ImageJ. Particles that remained stationary or underwent repetitive bidirectional movements for the majority of the recording time were defined as stationary/dynamic pause events. Each trajectory of stational/dynamic pause particles that could be visually separated was included in the number of stationary/dynamic pause events. The number of anterograde or retrograde events was counted in the same way. Stationary/dynamic pause percentage was calculated as the number of stationary/dynamic events divided by the sum of the number of anterograde, retrograde, and stationary/dynamics pause events. Anterograde and retrograde percentages were calculated similarly.

### Colocalization analysis

Control neurons were plated at a density of 1.8*10^3^ cells/mm^2^ on PDL-coated 12mm diameter coverslip in 24-well culture plate (about 3.5*10^5^ cells on 190 mm^2^ surface). Through transfection, pmCherry-n1-NMNAT2 was co-expressed with pEGFP-n1-APP, pEGFP-n1-SYPH, and pEGFP-c1-SNAP25 at DIV6 and samples were fixed using 4% PFA with 4% sucrose dissolved in PBS at DIV8. Immunocytochemistry was conducted to amplify the mCherry and EGFP signals after fixation. ProLong™ Gold antifade mounting medium with DAPI was used to mount the coverslips. Fluorescent confocal images were taken by a Leica TCS SPE confocal microscope using a 63x (1.4 N.A.) oil objective lens with system optimized z-step size. The colocalization was further validated by structure illumination imaging using OMX-SR 3D-SIM Super-Resolution System with a 60x (1.516 N.A.) oil objective lens, following system optimized z step size.

JACoP (Just Another Colocalization) imageJ plugin was used to analyze confocal images. Briefly, cell debris signal outside axon was manually deleted. “Objects based methods” was used. The threshold for both channels was manually adjusted based on the visual judgement of signal coverage. Minimum particle size was set as 25 pixel. The ratio of particles colocalized to NMNAT2 was calculated as the number of colocalizing particles in APP, SYPH, or SNAP25 channels divided by the total number of particles in these channels.

### Mitochondria density and morphology analysis

Both control and KO neurons were plated at a density of 1.8*10^3^ cells/mm^2^ on PDL-coated 12mm diameter coverslip in 24-well culture plate (about 3.5*10^5^ cells on 190 mm^2^ surface). Neurons were co-transfected with pCMV-MitoVenus and pCAG-mCherry at DIV6 and fixed using 4% PFA with 4% sucrose at DIV8. Immunocytochemistry with antibodies against GFP (Chicken) and RFP (Rabbit) was conducted to immuno-amplify the MitoVenus and mCherry signals after fixation. ProLong™ Gold antifade mounting medium with DAPI was used to mount the coverslips.

Images were taken by a Nikon A1 laser scanning confocal microscope using a 1.4 N.A. Apo Lambda S 60x oil objective lens with 0.171 μm z-step size and 3 times zoom to cover the whole depth of the axon segment of interest. Axonal segments at least 400 μm away from the soma were identified in the RFP channel and selected for imaging. Distal axons from at least 8-10 neurons were randomly sampled per coverslip and two coverslips were imaged per group. Images were analyzed using Imaris (Oxford Instruments). RFP channel was used to generate the surface masking of the axonal region, within which the MitoVenus signal was used to create the surface representing mitochondrial morphology. To calculate mitochondria density, length of the axon segment in each image was measured. Density was calculated from mitochondria number divided by axonal length in each image. Sphericity values of mitochondrial surface from axon segments of each individual neuron were input in each column in the column table in Prism 9.0 (each column representing one individual neuron). Cumulative frequency distribution of relative percentage was generated through “column analysis” function in Prism 9.0. Statistical analysis of cumulative frequency distribution is described in “quantification and statistical analysis” section.

### NAD^+^/NADH-Glo™ bioluminescent assay

Control or KO neurons were cultured in a 96-well plate at a density of 1.3*10^3^ cells/mm^2^ (4.235*10^4^ cells onto 32 mm^2^ surface). Following manufacturer’s instruction (Promega), neuronal cultures at DIV8 were washed three times with PBS and lysed in buffer containing 25 μl PBS and 25 μl 1 % DTAB (Dodecyltrimethylammonium bromide) base buffer (100 mM Sodium Carbonate, 20 mM Sodium Bicarbonate, 10 mM Nicotinamide, 0.05 % TritonX-100, pH 10-11). 30 μl of lysate was collected for BCA protein assay. 20 μl of the remaining lysate was diluted into 105 μl. 50 μl of diluted lysate was mixed with 25 μl 0.4 N HCl and incubated at 60°C for 15 min to detect NAD^+^; while another 50 μl of diluted lysate (without 0.4 N HCl) was incubated at 60°C for 15 min to detect NADH. After incubation, 25 μl of 0.5 M Tris base was added to the wells used for detecting NAD^+^ to neutralize the HCl; 50 μl of a solution containing 1:1 0.4 N HCl and 0.5 M Tris base was added to wells used for detecting NADH. Freshly dissolved NAD^+^ in PBS was used to prepare the standard curve. Luminescent signal from each sample was converted into absolute NAD^+^ or NADH concentration and normalized to protein concentration measured by the BCA method (Thermo Scientific™).

NAD^+^/NADH live imaging using the genetically encoded sensor, SoNar Around 1.66*10^3^ cells/mm^2^ cortical neurons were plated on PDL (Poly-D-Lysine)-coated MaTek dish (P35G-1.5-20-C). The NAD^+^/NADH sensor, SoNar, and its control cpYFP were gifts from Yi Yang’s group. On DIV6, 2.5 ug DNA of pCDNA3.1-SoNar or pCDNA3.1-cpYFP were transfected. After 48 h expression of the sensor or control (DIV8), the culture medium was replaced by Hibernate E buffer and maintained at 37°C for imaging using a Leica TCS SP8 confocal laser scanning platform with HyD hybrid detector and a HC PL APO 40x (1.2 NA) water-immersion objective. The SoNar sensor/cpYFP was excited at 440 nm and 488 nm and the emission at 530+/-15 nm was detected to obtain the ratiometric measurement. Laser power intensity was maintained the same between WT/Het and KO in the same experiment. Axonal segment > 400 μm away from the soma (defined as distal axon) and axonal segment within 200 μm to the soma (defined as proximal axon) were imaged with 1 μm z-step size to cover the whole axon segment within the field of view. Maximum intensity projection was generated for further analysis. Mean grey value from 488 nm excitation (F488) and 440 nm excitation (F440) was measured in ImageJ using manually drawn regions of interest (ROIs) that circle the edges of the axons of interest. Five rectangular ROIs were randomly drawn in the background area and the average mean grey value from these five ROIs was used to subtract background. After background subtraction, the ratio was calculated as (F488_axon_-F488_background_)/ (F440_axon_-F440_background_). Data presented were normalized to control as percentage.

### Syn-ATP live imaging

Syn-ATP imaging was performed using a custom-built inverted spinning disk confocal microscope (3i imaging systems; model CSU-W1) with two cameras: an Andor iXon Life 888 for confocal fluorescence imaging and an Andor iXon Ultra 888 electron-multiplying charge-coupled device camera for luminescence imaging. The Andor iXon Ultra 888 camera was selected for ultralow dark noise that was further reduced by cooling to −100°C. The speed of the Andor iXon Ultra 888 camera used for luminescence measurements was 1 MHz with 1.00 gain and 1000 intensification. Image acquisition was controlled by Slidebook 6 software. Confocal imaging of mCherry fluorescence was performed by illuminating neurons with a 561 nm laser with 200 ms exposure and 2.3 mW laser power. For mCherry fluorescence measurements, ten frames of timelapse imaging without interval were acquired and the average of ten images was used to measure the mCherry fluorescence signal from the varicosities of interest. For luminescence measurements, luminescence photons were collected by accumulating the image for 60 s in the presence of 2 mM D-luciferin (Promega), and the luminescence signal was measured from the same varicosities as the corresponding fluorescence image. All images were acquired through a Plan-Apochromat 63x/1.4 N.A. Oil objective, M27 with DIC III prism, using a CSU-W1 Dichroic for 561 nm excitation with Quad emitter and individual emitters for confocal fluorescence, and a 720 nm multiphoton short-pass emission filter was used for luminescence. During imaging, temperature was maintained at 37°C by means of an Okolab stage top incubator with temperature control. Distal axon at least 400 μm away from the soma was selected for imaging.

Images were analyzed in ImageJ using the plugin, Time series analyzer (https://imagej.nih.gov/ij/plugins/time-series.html). Regions of Interest (ROIs) of ∼1.2 μm diameter were drawn around the varicosities of interest to obtain the mean fluorescence and luminescence values from the corresponding fluorescence and luminescence images. We observed that some presynaptic varicosities in DIV8 neurons are not mature and not stable over the time course of imaging, and have a heterogenous amount of synaptophysin expression, consistent with previous studies [46]. Therefore, we focused on varicosities of at least 1.2 μm diameter and that were stable during the 60s luminescence acquisition. Neurons with an average mCherry fluorescence value lower than 1000 A.U. in their varicosities were excluded from the analysis, as low fluorescence value indicates low expression of the sensor and therefore resulted in low luminescence signal-to-noise ratio. For background subtraction of the fluorescence and luminescence values, three rectangular ROIs were drawn in the background region surrounding axons of interest. The average mean values from these background regions in the fluorescence (F_background_) and luminescence (L_background_) channels were used to subtract background for each individual varicosity from the fluorescence (F_varicosity_) and luminescence (L_varicosity_) channels, respectively.

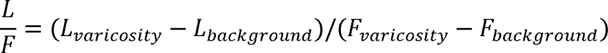

The average L/F value across varicosities within one neuron was used to represent the relative presynaptic ATP level for that neuron, accounting for one data point in the statistical analysis.

### pH measurement and pH correction of L/F measured from Syn-ATP

Cytoplasmic pHluorin (Cyto-pHluorin) was transfected in cortical neurons at DIV7 and imaging was done at DIV8 using the same equipment setting as Syn-ATP measurements. Detail procedures are described in the previous publication [33]. In brief, neuronal culture medium was replaced by Tyrodes buffer containing (in mM) 119 NaCl, 2.5 KCl, 2 CaCl_2_, 2 MgCl_2_, 30 HEPES (buffered to pH 7.4 at 37°C) and 25 glucose. Basal fluorescence intensity of Cyto-pHluorin was monitored for 5 min, acquired every 15 s (20 timepoints). ROIs were drawn around the axonal segment of interest and the mean fluorescence value was measured over the 20 timepoints (F_axon, frame i_). Three rectangular ROIs were drawn in the background region surrounding the axon to measure the average background value across the 20 timepoints (F_background, frame i_). F_basal_ was calculated by subtracting background in each timepoint and then averaging the value across the 20 timepoints.

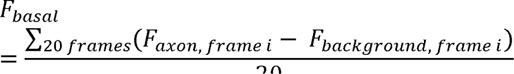

Then, 1 μM Oligomycin was added to the Tyrodes buffer and incubated for 5 min. Fluorescence of Cyto-pHluorin was imaged to obtain F_Oligo_ using the above calculation.

Then, Tyrodes buffer was replaced by an NH_4_Cl solution containing (in mM) 50 NH_4_Cl, 70 NaCl, 2.5 KCl, 2 CaCl_2_, 2 MgCl_2_, 30 HEPES (buffered to pH 7.4 at 37 °C) and 25 glucose. The response of Cyto-pHluorin to NH_4_Cl treatment was monitored for 5 min, every 15 s (20 timepoints). The measured fluorescence of Cyto-pHluorin was called F_Max,_ and was background subtracted as explained above.

Cytosolic pH of the basal and Oligomycin treatments were determined using the following modified Henderson-Hasselbalch equation:

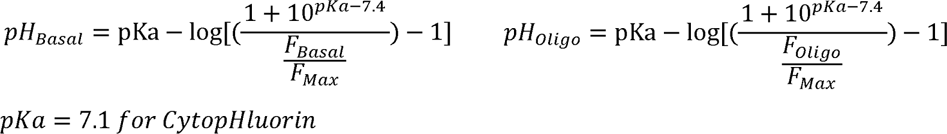

From the known pH values at various conditions, Syn-ATP L/F value was corrected as follows:

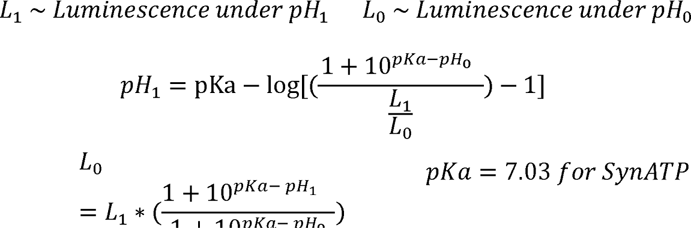

Here, the average pH in each condition was used for correction and we did not propagate errors of pH measurements into the final error shown in L/F measurements. pH corrections of L/F values were only done for those conditions that showed a statistically significant change in pH compared to control (as in Fig 5-S1).

### NAD^+^ supplementation in neuronal culture

NAD^+^ (Roche, NAD100-RO) was dissolved in PBS at a stock concentration of 100 mM, sterilized, aliquoted, and stored at -80°C. Only less than 1-week-old stocks were used. At the starting day of supplementation, NAD^+^ was applied to the culture medium at a final concentration of 1 mM. In the following days before live imaging, 1/3 of culture medium was replaced with fresh medium containing 1 mM NAD^+^ every day. 1 mM NAD^+^ was supplemented to the Hibernate E low fluorescence buffer used for live imaging. In the following days before fixation, 1/3 of culture medium was replaced with fresh medium containing 1 mM NAD^+^ every 3 days.

### 2->*Deoxyglucose* (2DG) treatment in neuronal culture

To acutely inhibit glycolysis while leaving mitochondrial respiration intact, 15 mM 2-DG and 10mM methyl-pyruvate were added to the customized Hibernate E low fluorescence buffer containing 0 mM glucose (∼262 mmol/kg osmolality). Culture medium was replaced by the 2DG Hibernate E buffer and neurons were incubated for 30 min before imaging APP-EGFP axonal transport. For each culture dish, imaging of axonal transport was conducted within 30 min of 2DG treatment. To chronically inhibit glycolysis, 25 mM 2-DG and 10 mM methyl-pyruvate were added to the neurobasal medium containing 25 mM glucose. 2 days after treatment, neuronal culture was fixed for immunocytochemistry.

### Oligomycin treatment in neuronal culture

To acutely inhibit ATP synthesized from mitochondrial respiration, 1 μM oligomycin was added to the commercial Hibernate E low fluorescence buffer containing 25 mM glucose (∼241 mmol/kg osmolality). Culture medium was replaced by Oligo Hibernate E buffer and neurons were incubated for 5 min before imaging APP-EGFP axonal transport, Syn-ATP or Cyto-pHluorin. For each culture dish, live imaging was conducted within 30 min of oligomycin treatment. For APP axonal transport imaging, oligomycin (Calbiochem 495455) was dissolved in DMSO to prepare a 1 mM stock. For Syn-ATP and Cyto-pHluorin imaging, oligomycin (Sigma-Aldrich O4876) was dissolved in 100% Ethanol to prepare a 1 mM stock.

### Antisense Oligonucleotide (ASO) treatment in neuronal culture

The antisense oligonucleotides (ASOs) were designed, synthesized, and preliminarily tested by Ionis Pharmaceuticals. ASOs consist of 20 chemically modified nucleotides with a central gap of 10 deoxynucleotides flanked on its 5′ and 3′ sides by five 2′-O-(2-methoxyethyl) (MOE)-modified nucleotides. Thirteen of the phosphodiester internucleotide linkages were replaced with phosphorothioate to further enhance the stability of ASOs to endogenous nucleases. The non-targeting ASO, 5’-CCTATAGGACTATCCAGGAA-3’ (ASO30), was used as control. The ASOs targeting mouse SARM1 mRNA, 5’-GGTAAGAGCCTTAGGCACGC-3’ (ASO33) and 5’-CCACCTTTTAGTCAAGACCC-3’ (ASO47) were applied to silence SARM1 expression. Both ASOs arrived as 100mg/ml stock. At the treatment starting day as indicated, ASOs were diluted into 5µM as the final concentration in neuronal culture medium. In the following days before live imaging or fixation, 1/3 of culture medium was replaced with fresh medium containing 5 µM ASOs every 3 days.

### Quantitative real-time PCR

Following ASO treatment, total RNA was extracted from neuronal culture by RNeasy Mini Kit (Qiagen 74104) and followed by on column DNase digestion according to the manufacturer’s instruction. RNA concentration and purity was measured by NanoDrop 2000 spectrophotometer (ThermoFisher Scientific). 500ng RNA was used as input for cDNA synthesis through iScript™ advanced cDNA synthesis kit (Bio-Rad 1725038) following the manufacturer’s protocol. To quantify mRNA level, quantitative real-time PCR was conducted using PrimeTime™ Gene Expression Master Mix (IDT 1055772) following the manufacturer’s instruction.

To detect SARM1 mRNA, 0.3 ul of cDNA was loaded into the reaction mix (10 ul in total) containing 450 nM of SARM1 forward primer, 450 nM of SARM1 reverse primer, and 150 nM SARM1 probe. Mouse SARM1 primer probe set was customized through PrimeTime™ qPCR Probe Assays (IDT). SARM1 forward primer sequence is 5’ TTTGTCCTGGTGCTGTCTG 3’. SARM1 reverse primer sequence is 5’ GCCACTCAAAGCCATCAATG 3’. SARM1 probe sequence is 5’ ACAATCTCCTTGTGCACCCAGTCC 3’, with FAM reporter at 5’, internal ZEN quencher, and 3’ Iowa Black® FQ quencher. To detect GAPDH mRNA as the internal control for normalization, 0.3 ul of cDNA was loaded into the reaction mix (10 ul in total) containing 500 nM of GAPDH forward primer, 500 nM of GAPDH reverse primer, and 250 nM GAPDH probe. Mouse GAPDH primer probe set was acquired from PrimeTime™ Predesigned qPCR Assays with same reporter and quencher (Assay ID: Mm.PT.39a.1). These assays were run in QuantStudio 7 Flex Real-Time PCR Systems (ThermoFisher Scientific). The Ct value was defined as the number of cycles required for the fluorescence to exceed the detection threshold. The ΔCt values were calculated by subtracting Ct value of GAPDH from SARM1. The ΔΔCt values were calculated by subtracting the average ΔCt of non-targeting ASO treated control from the ΔCt of each individual sample. The relative fold change was calculated as 2^−ΔΔCt^.

### Western blotting

Following ASO treatment, neuronal culture was lysed in RIPA buffer [20 mM of Tris-base, 150 mM of NaCl, 2 mM EDTA, 1% Triton X-100, 0.5% sodium deoxycholate, 0.1% sodium dodecyl sulfate (SDS), 1x cOmplete™ protease inhibitor cocktail and phosphatase inhibitor cocktail, pH 7.4], and homogenized by motorized disposable pellet mixers for 15 s, then sonicated by VMR Branson Sonifier 250 10 times under the parameter: duty cycle 30 and output 3.

Protein concentration was measured using the BCA method (Thermo Scientific™). 10-15 ug proteins were separated on a 10% SDS-polyacrylamide gel and then transferred to a nitrocellulose membrane of 0.45 μm pore size (BioRad). To detect SARM1 protein level, the membranes were incubated overnight with the homemade primary antibody against SARM1, which was a gift from Dr. Yi-Ping Hsueh [47]. Next day, the membranes were incubated with a species-appropriate secondary antibody. Western blot images were acquired by LI-COR Odyssey scanner and software (LI-COR Biosciences, Lincoln, NE USA) and quantified with NIH ImageJ software.

### TUJ1 fragmentation quantification

Fluorescent confocal images of TUJ1 staining were taken in the way described in "immunocytochemistry and confocal imaging” section. TUJ1 fragmentation was analyzed following the degeneration index measurements method described by Kraemer et al. [48], with some modifications to their particle size and circularity indexes. TUJ1 fragmentation quantification was performed using ImageJ. The fluorescent channel corresponding to TUJ1 was isolated and converted to a binary image using the “Make Binary” function and this image was used for all subsequent steps of the analysis. Total TUJ1 expression area was measured using the “Measure” function. Images were then processed using the “Analyze Particles” function, setting the size parameter at 0.327 µm^2^-817.587 µm^2^ (4-10000 pixels), and circularity at 0.2-1. The area of selected particles was recorded and normalized to total area of TUJ1.

### Experimental Design

No statistical methods were used to predetermine the sample size. For in vivo studies, investigators were blinded to the mouse genotype during data acquisition and analysis. For in vitro manipulation, investigators were not blinded to the genotype of neuronal cultures and treatments. During the quantification of APP accumulation area, TUJ1 area, MAP2 area, TUJ1 fragmentation degree, and the analysis of axonal transport, investigators were blinded to the genotype and treatment information of each image. For NAD/NADH-Glo™ bioluminescent assay, investigators were not blinded, and quantification was done by plate-reader without human curation. During the acquisition and quantification of NAD^+^/NADH (SoNar) sensor imaging, Syn-ATP sensor imaging, and Cyto-pHluorin imaging, the investigators were not blinded to the genotype and treatment of each image. Inclusion and exclusion criteria applied during the analysis of Syn-ATP sensor imaging to avoid varicosities with extremely low mCherry (indicator of sensor copy number) expression. For Western Blotting and qPCR, investigators were not blinded to genotype and treatment. Except gross neurite area measurement at DIV5 & DIV14 and Cyto-pHluorin live imaging, all experiments were replicated with culture from at least 2 independent primary culture preparations, as detailed in the supplementary excel sheets.

### Quantification/statistical analysis and figure constructions

Images were analyzed using Fiji (Image J with updated plug-ins) and Imaris. NAD/NADH-Glo™ bioluminescent Assay were measured by CLARIOstar plate reader. GraphPad Prism 9.0 (GraphPad Software) was used for statistical analysis. The outliers were detected by “ROUT” method in GraphPad Prism 9.0 and excluded for statistical analysis. The normality of residuals was mostly checked by D’Agostino-Pearson normality test as recommended by the GraphPad Prism user guide. For data with normally distributed residuals, one-way ANOVA, two-way ANOVA, or unpaired student’s t-test was used. For data that did not pass the D’Agostino-Pearson test, one-way ANOVA was replaced by Kruskal-Wallis test; two-way ANOVA was replaced by Multiple Mann-Whitney test or re-formatted for Kruskal-Wallis test; unpaired student’s t-test was replaced by the Mann-Whitney test.

To analyze cumulative frequency distribution curve, the dependent variable P (the relative percentage within each bin generated by GraphPad Prism “Column analysis”) was transformed into log(P/(100-P)) so that the relationship between covariate (bin of sphericity) and the transformed dependent variable was linearized. Then, a linear mixed model with random slope and intercept was generated by SPSS using a restricted maximum likelihood approach to compare statistical difference between control and KO. Criterion for statistical significance was set at p < 0.05 for all statistical analyses.

Figures were made with Adobe Photoshop CS6 and Illustrator CS6 and brightness/contrast, orientations, and background corrections were applied to better illustrate the staining patterns.

## Results

### Deleting NMNAT2 in post-mitotic cortical glutamatergic neurons results in deformed brains and age-dependent loss of long-range cortical axons

Germline deletion of NMNAT2 in mice results in premature death at birth with severe axonal outgrowth deficits and subsequent degeneration in peripheral and optic nerves [37, 39]. Using explant cultures prepared from NMNAT2 KO embryonic cortices, Gilley et al. showed NMNAT2 KO cortical axons were shorter than control axons and exhibited degenerative phenotypes [37]. The premature death of germline NMNAT2 KO prevented in vivo examinations of the role of NMNAT2 in cortical neurons. Using mRNA in situ hybridization technique, we found that nmnat2 mRNA is mainly enriched in the embryonic cortical plate, where post-mitotic glutamatergic neurons are located (Fig1-S1A). Axons of cortical glutamatergic neurons often travel long distances to the contralateral hemisphere or to subcortical regions via extensive and complex arbors [6]. These observations led us to hypothesize that NMNAT2 is required for axonal outgrowth and the health of cortical glutamatergic axons.

To test this hypothesis, we generated NMNAT2 conditional knockout mice (cKO) by crossing NMNAT2 conditional allele mice (NMNAT2^f/f^) [37] with a Nex-Cre transgenic mouse line, which expresses Cre recombinase in post-mitotic glutamatergic neurons mainly in the cortex and hippocampus from embryonic day 11.5 (E11.5) [38] (Fig1-S1B). Despite the restricted deletion of NMNAT2, only about 50% of cKO mice survived after birth, with no apparent lethality at E18.5 (Fig1-S1C-D). The surviving cKO mice weighed significantly less than their littermate controls from early postnatal ages through adulthood (Fig. 1A). NMNAT2 cKO mice exhibited evident hindlimb clasping (Fig. 1B), ataxia and forelimb circling phenotypes (10 out of 10 mice examined), reflecting motor behavioral deficits similar to those observed in many neurodegenerative mouse models [49–51]. cKO brains were visibly smaller than controls from postnatal ages on (Fig. 1C).

**Fig 1.**
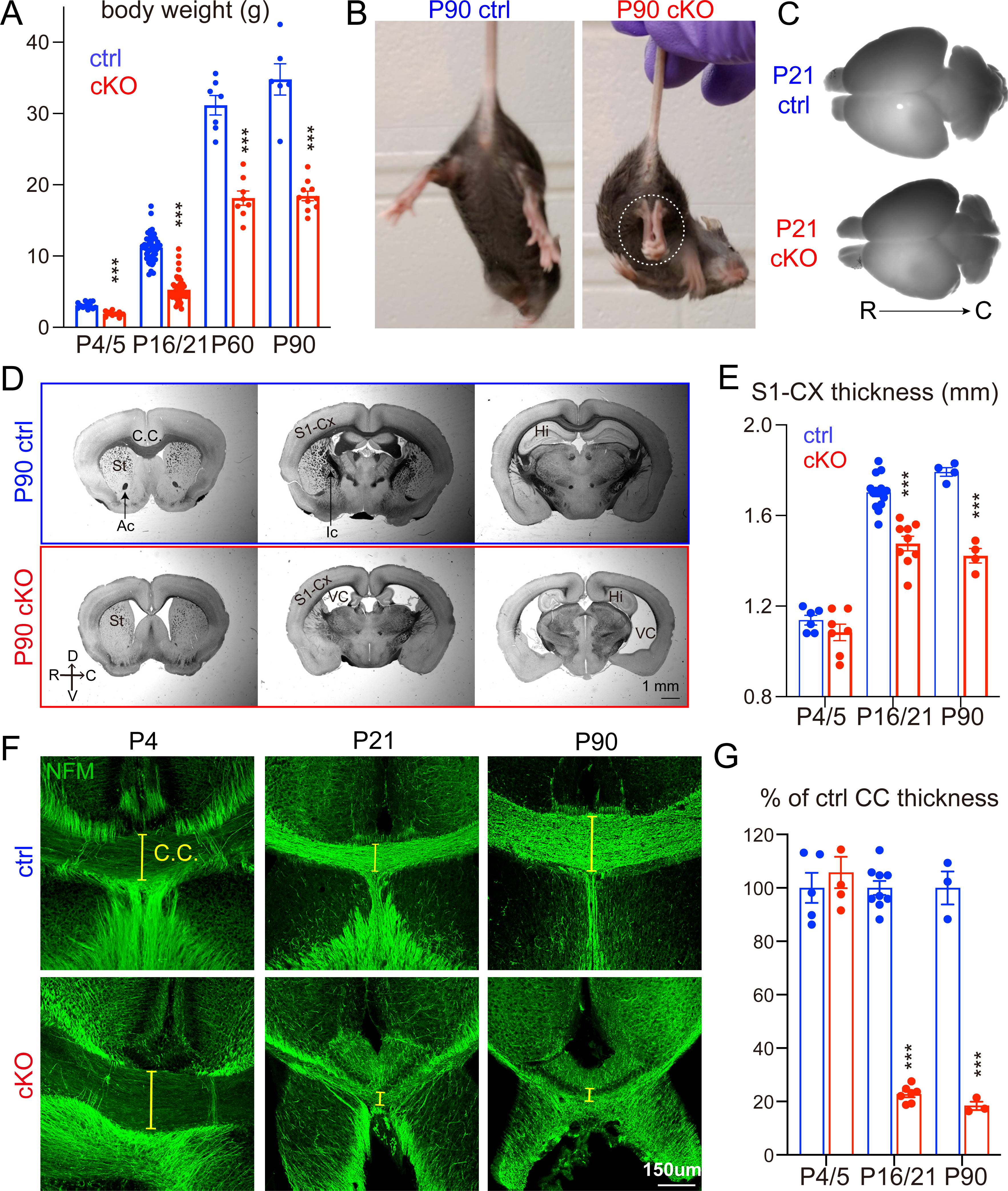
Deleting NMNAT2 in cortical glutamatergic neurons results in age-dependent axonal degeneration. **(A)** Body weight of cKO and their littermate control (ctrl) mice at P4, P16/21, P60, and P90. Mice numbers: P4/5, 13 ctrl and 9 cKO; P16/P21 43 ctrl and 46 cKO; P60, 7 ctrl and 8 cKO; P90, 6 ctrl and 10 cKO. **(B)** Movie screenshots showing that a P90 cKO mouse exhibits hindlimb clasping behaviors (dashed white oval), a classic motor deficit observed in many neurodegenerative models, but not in a ctrl mouse (see Sup. Movies). **(C, D)** Bright field images showing whole brains and coronal plane brain sections (rostral to caudal from left to right) from ctrl and cKO mice. In addition to the smaller brain sizes, cKO brains have enlarged ventricles and reduced cortical regions and hippocampal areas. **(E)** Quantification of the primary somatosensory (S1) cortex thickness in ctrl and cKO mice at different ages. Mice numbers: P4/5, 6 ctrl and 7 cKO; P16/P21, 14 ctrl and 9 cKO; P90, 4 ctrl and 4 cKO. **(F)** Confocal images of immunohistochemical staining of NFM (medium-size neurofilament) showing axonal tracts through the corpus callosum (CC) in ctrl and cKO brains at P4, P21, and P90. Yellow brackets mark the thickness of the CC. **(G)** Quantification of the CC thickness, normalized to its value in ctrl mice. Mice numbers: P4/5, 5 ctrl and 5 cKO; P16/P21, 9 ctrl and 7 cKO; P90, 3 ctrl and 3 cKO. Abbreviations: Ac, anterior commissure; Ic, internal capsule; CC, corpus callosum; Cx, cortex; Hi, hippocampus; St, striatum; VC, ventricle. Unpaired t-test and Mann-Whitney test were applied for the statistic result, ***p<0.001, ****p<0.0001.

To evaluate gross brain morphology, we examined brain sections along rostral to caudal coronal planes with bright field microscopy. Compared to the brains of littermate controls, the brains of cKO mice displayed enlarged ventricles, smaller hippocampi, aberrant anterior commissures, a thinner corpus callosum, and a thinner cortex (Fig. 1D). We measured the thickness of the primary somatosensory cortex and found that it was significantly reduced in cKO compared to their littermate controls at postnatal day 16/21 (P16/21) and P90 (Fig. 1E). The degenerative brain phenotypes observed in NMNAT2 cKO brains highlight the importance of NMNAT2 in cortical neuronal health.

To distinguish whether brain dystrophy in NMNAT2 cKO is caused by axonal outgrowth deficits versus axonal maintenance failures, we examined the corpus callosum, where the long-range callosal axons of cortical glutamatergic neurons cross the midline on the way to their contralateral targets [8, 9]. To facilitate visualization of the corpus callosum, we immunostained the medium-size neurofilament (NF-M), a cytoskeleton protein enriched in axons, in P4/5, P16/21, and P90 cKO and control brains. There was no difference in corpus callosum thickness between control and cKO brains at P4/5 (Fig. 1F-G). However, a drastic reduction of corpus callosum thickness in cKO mice occurred by P16/21 and persisted until at least P90, the eldest age examined (Fig. 1F-G). Most callosal axons have already finished midline crossing at P4/5 [52, 53]. Thus, the normal corpus callosum thickness in cKO mice at P4/5 suggests that the deletion of NMNAT2 in glutamatergic neurons does not impair initial axonal outgrowth. Instead, the age-dependent reduction of corpus callosum thickness and degeneration-like brain morphology suggest that NMNAT2 is required to maintain the health of long-range cortical axons.

### NMNAT2 loss leads to APP accumulation in axons prior to degeneration *in vivo and in vitro*

Based on the axon degeneration phenotype in NMNAT2 cKO brains, we hypothesized that NMNAT2 loss disrupts axonal physiology, ultimately resulting in axonal degeneration. Axonal transport plays a critical role in neuronal function and survival [28] and its deficits are thought to be a primary cause of axonopathy [54]. Amyloid Precursor Protein (APP) is rapidly and bidirectionally transported in axons [55–59] and is required for synaptogenesis and synaptic function, plasticity, etc. [60–62]. Axonal APP accumulation has been found in AD mouse models [63, 64] and traumatic brain injury [65, 66], and serves as a marker for axonal transport breakdown [67].

To examine whether NMNAT2 deletion results in axonal transport deficits, we immunostained APP in brains prepared from P5 and P21 cKO mice and littermate controls. We found significant APP accumulation in corpus callosum at P5 in cKO but not in control (Fig. 2A,E). This is before the corpus callosum thickness is affected by NMNAT2 loss. In addition, we observed APP accumulation in cKO mice in brain regions where glutamatergic axons transit, including the hippocampal fimbria and striatum. In contrast, little APP signal was detected in the corresponding regions of control brains (Fig. 2B,C,E). At P21, where significant axon degeneration occurs, we found a striking buildup of the APP signal in areas enriched with long-range axons. The observation of APP accumulations encapsulated by myelin basic protein (MBP), a marker for myelinated axons, in the corpus callosum (Fig. 2D), suggests that the APP aggregates detected are present in axons.

**Fig 2.**
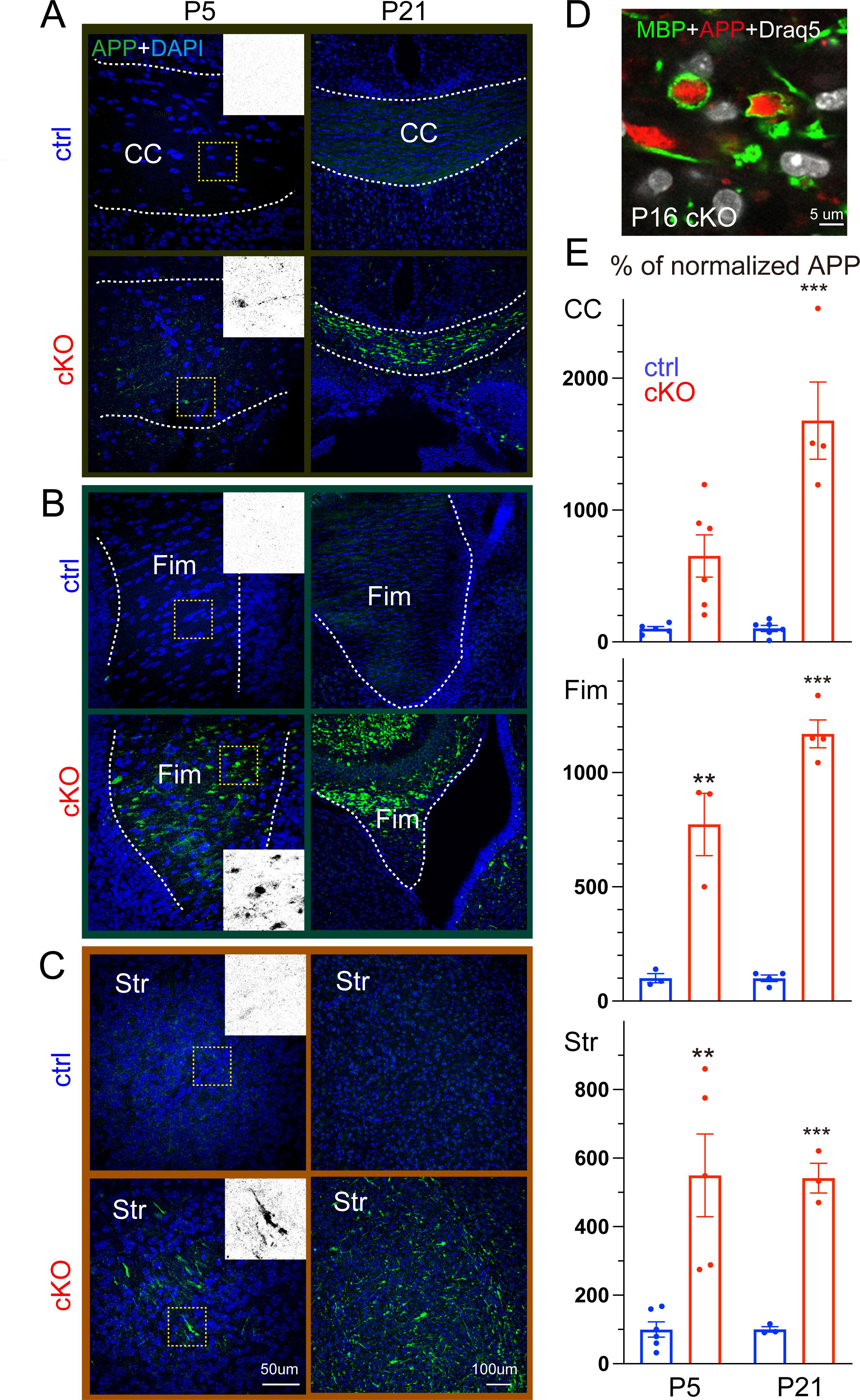
Deleting NMNAT2 in cortical glutamatergic neurons results in Amyloid Precursor Protein (APP) accumulation. **(A, B, C)** Representative confocal microscopy images showing APP accumulation in multiple brain regions of ctrl and cKO mice at P5 and P21. Scale bars, 50 um for all left panels, 100 um for all right panels. Dashed white lines mark the margins of corpus callosum (A) and fimbria (B). **(D)** High magnification image showing that the accumulated APP is encircled by myelin sheath labeled by myelin basic protein (MBP) staining in P16 cKO corpus callosum (observed in 3 out of 3 cKO brains). (E) Quantification of the APP signal in the corpus callosum, hippocampal fimbria and striatum area of cKO mice, normalized to the average signal in ctrl mice, at P5 and P21. Number of mice: P5, 3-6 ctrl, and 3-6 cKO; P16/P21 3-6 ctrl and 3-4 cKO. Unpaired t-test and Mann-Whitney test were applied for the statistic result, *p<0.05, **p<0.01, ***p<0.001.

Interestingly, we also found analogous APP accumulations in the neurites of cortical neuron cultures prepared from NMNAT2 null (NMNAT2-Blad [39]) embryos, but not in wildtype or heterozygous (control) neurons after 8 days in vitro (DIV; Fig. 2-S1&S2). Immunostaining showed APP accumulation in axons labeled by βIII-tubulin/TUJ1 antibody but not in MAP2-positive dendrites of KO neurons (Fig. 2-S1C,D). APP accumulation increased drastically in KO axons from DIV8 to 14 (Fig. 2-S1A, B), while the signal densities for TUJ1 and MAP2 levels were similar between KO and control neurons (Fig. 2-S2A,B). Additionally, we detected fragmented and aggregated TUJ1 signal as a sign of axon degeneration at DIV14 (Fig. 2-S1 and Fig. 8-S2C). As our cortical neuronal cultures are prepared from E15.5/E16.5 embryonic cortex, by DIV8 they likely correspond to the first postnatal week of age in vivo. Additionally, we found significant axonal APP accumulation in NMNAT2-deleted cortical neurons using an alternative Cre-loxP approach (Fig. 2-S3). This in vitro recapitulation of APP accumulation and axon degeneration-like phenotype seen in vivo justifies the use of cultured NMNAT2 KO neurons as a cellular model to elucidate the molecular mechanisms mediating NMNAT2’s function in axonal health.

### NMNAT2 is required for the transport of fast-moving vesicular cargos in distal axons

As APP accumulation is observed prior to axonal degeneration in NMNAT2 KO axons, we hypothesized that NMANT2 is required for axonal transport. Previous work showed that NMNAT2 is localized to Golgi-derived and undergoes fast, bi-directional axonal transport in PNS axons [25, 26]. We observed a similar transport of NMNAT2 up and down axons and its colocalization and comigration with fast-moving cargos in cultured cortical neurons (Fig. 3-S1).

To determine whether NMNAT2 is required for axonal transport, we used time-lapse imaging to quantify axonal transport of EGFP-tagged vesicular cargos, APP and SNAP25, and DsRed-tagged mitochondria in control and NMNAT2 KO neurons at DIV6 and 8 (Fig. 3 and Fig. 3-S2). APP is a component of Rab5-containing vesicles [68], while SNAP25 is a component of Piccolo-bassoon transport vesicles [69]. Rab5 and SNAP25 undergo fast, bidirectional axonal transport [70, 71] mediated by kinesin-1 and dynein motor proteins [55, 72]. Distinct from vesicular cargos, axonal mitochondria are known to move intermittently in both directions and more slowly [73, 74], in response to the interplay between adaptor and motor proteins [75, 76]. To transfect only a few neurons with APP-EGFP, SNAP25-EGFP, or mito-DsRed expressing construct, lipofectamine transfection was conducted <20 h before live imaging to reduce toxicity of overexpression. Such sparse labeling allowed us to identify distal (>400 μm away from the soma) or proximal (within 200 μm of the soma) axon segments (Fig. 3-S3). Axonal transport of APP and SNAP25 was measured in distal and proximal segments (Fig. 3 and Fig. 3-S2). At DIV8, significant deficits in APP and SNAP25-transport were detected in KO distal segments (Fig. 3B-G) but not proximal segments (Fig. 3-S2). Furthermore, we observed a substantial increase in the percentage of vesicles in the stationary and dynamic pause phases (Fig. 3C,F), a concomitant decrease in the percentage of vesicles engaging in anterograde movement, and reduced velocities in anterograde and retrograde directions (Fig. 3D, G). However, at DIV4 (not shown) and 6 (Fig. 3C,D,F,G), APP and SNAP25 transport were unaffected in KO axons. In contrast to vesicular transport, mitochondrial distribution, morphology, and motility were unaffected in KO neurons at DIV8 (Fig. 3-S4), despite the heterogeneous mitochondrial velocities reported previously [77]. These results demonstrate that NMNAT2 is required for fast transport of vesicular cargos but not mitochondria, and that NMNAT2 loss affects distal axon transport before axon degeneration.

**Fig 3.**
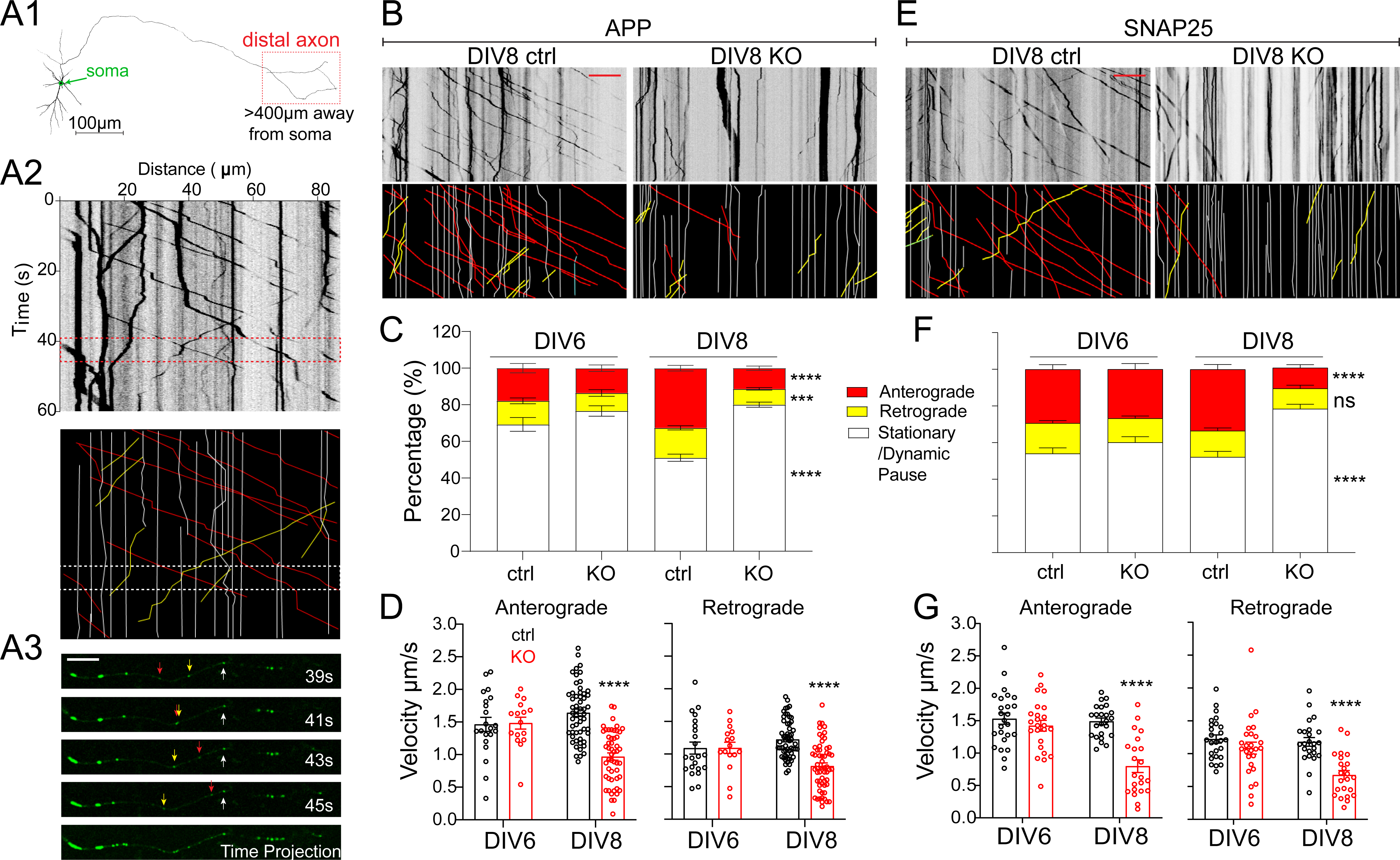
Nmnat2 is required for transporting vesicular cargos in the distal axon. **(A1)** Diagram indicating the distal axon region (> 400 µm from the soma) where axonal transport was examined with time-lapse imaging. **(A2)** Example kymograph (upper) and annotations (lower) with anterograde (red lines), retrograde (yellow lines), and stationary/dynamic pause (white lines) movements indicated. **(A3)** Four image examples showing APP-EGFP movement along an axon segment during the times indicated in the dashed box in **(A2)**. Scale bar, 10 µm. **(B, E)** Representative kymographs for APP-EGFP and SNAP25-EGFP from DIV8 ctrl and KO distal axons. Scale bar, 20 µm. **(C, F)** Percentages of APP-or SNAP25-EGFP vesicles moving antero/retrogradely or staying stationary/dynamic pause in ctrl and KO neurons. Numbers (neurons imaged) and statistics: **(C)** DIV6, 22 ctrl, and 19 KO; DIV8, 56 ctrl, and 62 KO; compared with two-way ANOVA and Šídák’s multiple comparisons for anterograde and stationary/dynamic pause categories, compared with multiple Mann-Whitney test and Holm-Šídák multiple comparisons for the retrograde category. **(F)** DIV6, 27 ctrl, 26 KO; DIV8, 24 ctrl, 27 KO; two-way ANOVA with Tukey’s multiple comparisons. **(D, G)** Velocity of antero-and retrograde transport in ctrl and KO neurons from 2-3 independent experiments. Numbers (neurons imaged) and statistics: **(D)** Anterograde velocity analysis: DIV6, 21 ctrl and 16 KO; DIV8, 56 ctrl and 51 KO. Retrograde velocity analysis: DIV6: 22 ctrl, 16 KO; DIV8, 56 ctrl, 58 KO. Two-way ANOVA with Šídák’s multiple comparisons. **(G)** Anterograde velocity analysis: DIV6, 25 ctrl, 24 KO; DIV8, 24 ctrl, 22 KO; two-way ANOVA with Tukey’s multiple comparisons. Retrograde velocity analysis: DIV6, 25 ctrl, 26 KO; DIV8, 24 ctrl, 23 KO; multiple Mann-Whitney test with Holm-Šídák multiple comparisons. KO was compared to ctrl of the same DIV in all data sets. Data represent mean ± SEM, ****p<0.0001. APP and SNAP25 data were collected from 3 and 2 independent experiments, respectively.

### NMNAT2 maintains global NAD, NADH levels, and local NAD /NADH redox potential in distal axons

NMNAT2 catalyzes NAD synthesis in the salvage pathway, the major pathway in CNS neurons for NAD biosynthesis [15]. The ratio of oxidized (NAD^+^) to reduced (NADH) forms of NAD establishes the NAD redox potential (NAD^+^/NADH) and is crucial to driving glucose metabolism [78, 79]. To determine if NMNAT2 is required for maintaining the NAD redox potential, we measured the abundance of NAD^+^ and NADH at DIV8, when axonal transport deficit first becomes evident in KO axons (Fig. 2-S1 C,D). Both NAD^+^ and NADH levels were reduced to ∼50% of their control value in KO neurons, suggesting that NMNAT2 is a major source of NAD in cortical neurons. However, since NAD^+^ and NADH levels were reduced to the same extent upon NMNAT2 loss, the NAD redox potential measured from whole neurons remained unchanged (Fig. 4A).

**Fig 4.**
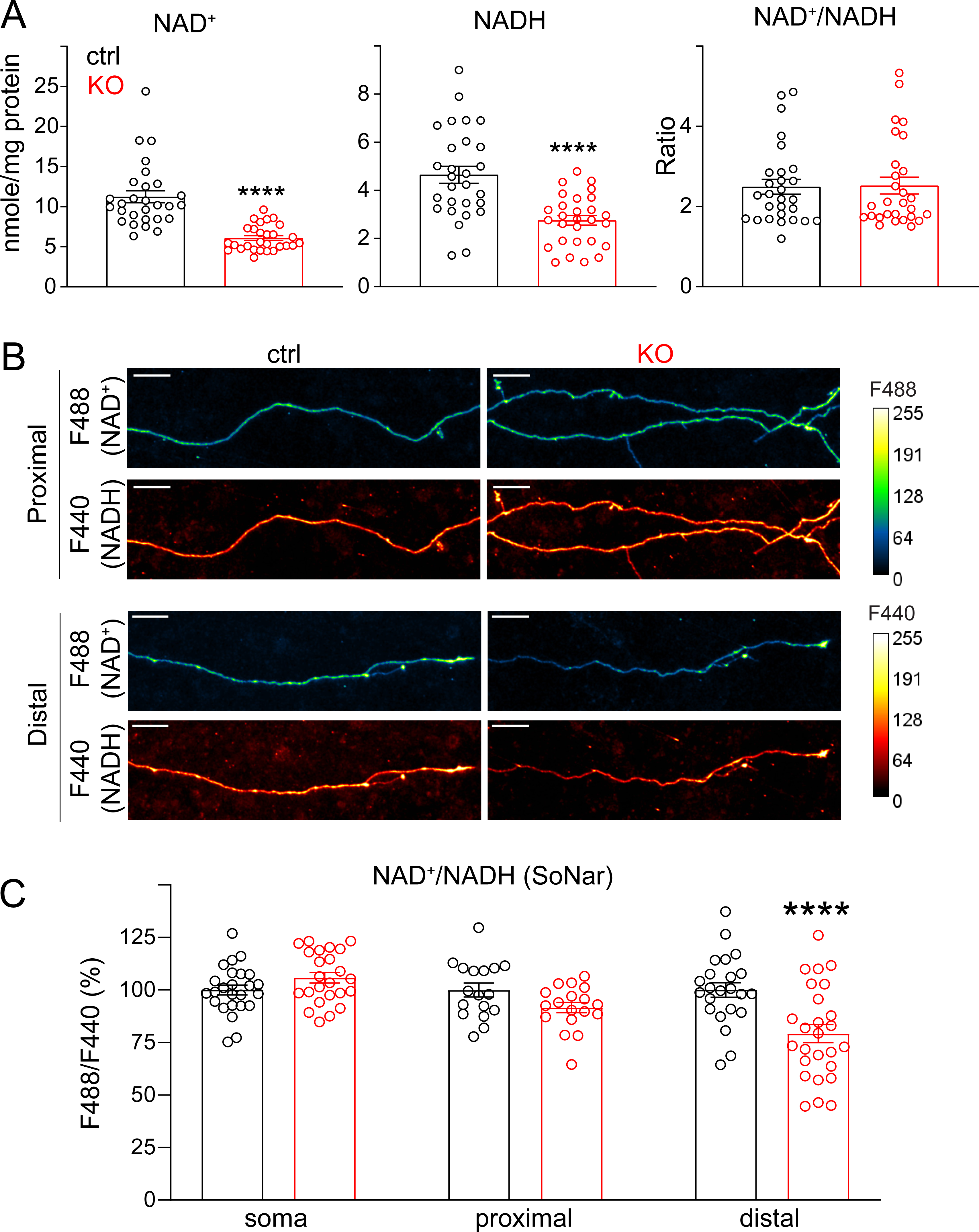
Loss of NMNAT2 reduces NAD levels and impairs the NAD redox potential in distal axons. **(A)** Levels of NAD^+^ and NADH and calculated NAD^+^/NADH ratios in DIV8 ctrl and KO cortical neurons as measured by the NAD^+^/NADH-Glo assay and normalized to protein amounts. Readings from 28 ctrl and 28 KO culture wells from 4 independent culture experiments; unpaired t-test was used for NADH while Mann-Whitney test was used for both NAD^+^ and NAD^+^/NADH ratios due to their distributions. **(B)** Representative images showing signals emitted from the SoNar (NAD^+^/NADH) ratiometric sensor for NAD^+^ and NADH in the proximal and distal axons of DIV 8 ctrl or KO cortical neurons. Scale bar, 20 um. **(C)** NAD+/NADH ratios in soma, proximal and distal axons of DIV7 and 8 ctrl and KO cortical neurons revealed by F488/F440 ratiometric measurements of the SoNar sensor (Soma: 25 ctrl and 24 KO from 2 independent experiments; proximal axon segments: 19 ctrl and 20 KO from 3 independent experiments; distal axon segments: 26 ctrl and 30 KO from 3 independent experiments; two-way ANOVA with Tukey’s multiple comparisons test). All above data represents mean ± SEM, ****p<0.0001.

One reason that NMNAT2 loss affects axonal transport in distal but not proximal axons may be that its homolog NMNAT1, which is expressed in the nuclei of neurons, is sufficient to maintain NAD redox potential in the soma and proximal axons [14, 80]. Thus, we hypothesized that NMNAT2 is only critical for maintaining NAD redox potential in distal axons, where it provides the ATP required for fast axonal transport. To measure NAD redox potential, we used a genetically encoded sensor, SoNar [44], that detects cytosolic NAD redox potential with a reliable signal-to-noise ratio and high spatial resolution. Our imaging studies found that NAD redox potential was significantly reduced in distal axons but not in the soma or proximal axons of DIV8 KO neurons compared to controls (Fig. 4B-C). These data confirm a subcellular requirement for NMNAT2 to maintain NAD redox potential in distal axons. In addition, they suggest that NMNAT2 contributes to ∼50% of the overall NAD^+^ and NADH levels in cortical neurons.

### NMNAT2 is critical for glycolysis on synaptic vesicles

Recent studies show that fast axonal transport of vesicular cargo is mainly fueled by on-board glycolysis through vesicle-associated glycolytic enzymes that generate ATP to fuel motor protein movement [32, 81]. Considering the transport deficits and the reduced NAD^+^/NADH redox potential in the distal axons of NMNAT2 KO neurons, we hypothesized that NMNAT2 supports fast axonal transport by facilitating local glycolysis. To measure ATP near fast-moving vesicular cargos, we transfected the genetically encoded presynaptic ATP sensor (Syn-ATP) in control and NMNAT2 KO neurons and measured sensor signals in distal axons at DIV 8 with live imaging (Fig. 5). This sensor is targeted to synaptophysin, one of the well-known fast vesicular cargos. Syn-ATP comprises an optimized luciferase to detect ATP by luminescence and an mCherry fluorescent protein as an internal control for sensor expression level [33] (Fig. 5B). The luminescence to fluorescence ratio (L/F) measured from Syn-ATP is proportional to ATP levels near synaptic vesicles (sv-ATP) [33]. As the Syn-ATP sensor is pH sensitive [33], all the measurements were pH-corrected (Fig. 5-S1A-C).

**Fig 5.**
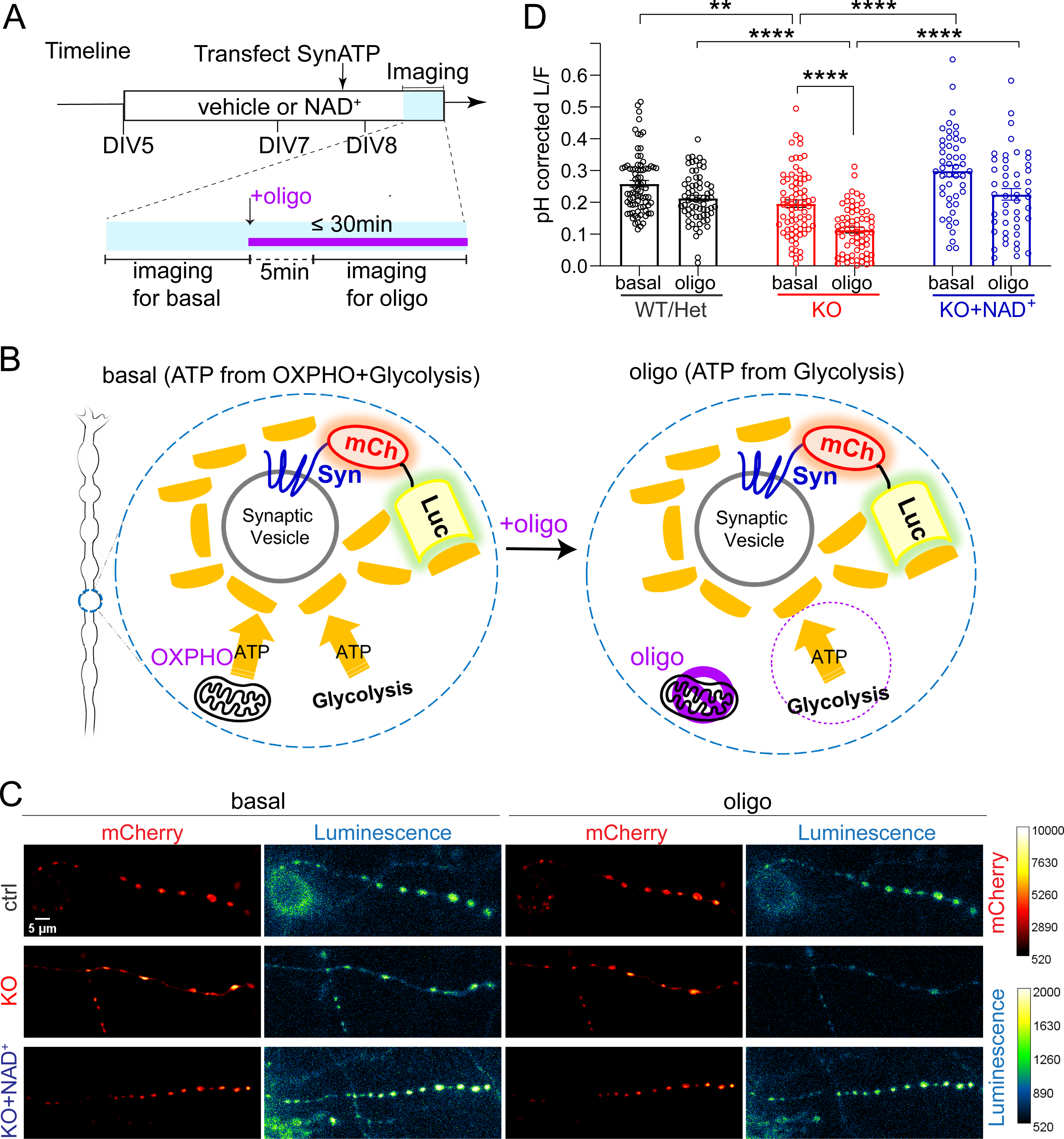
NMNAT2 loss impairs glycolysis in distal axon presynaptic varicosities. **(A)** Schematic diagram of the Syn-ATP sensor configuration and the treatment and imaging timeline. Syn: synaptophysin, mCh: mCherry, Luc: luciferase. **(B)** Representative Syn-ATP luminescence and fluorescence images of distal axon varicosities under basal and oligomycin-treatment conditions from DIV8 ctrl, KO, and KO neurons supplemented with NAD^+^ (KO+NAD^+^). Scale bar, 5 μm. **(C)** pH-corrected L/F (Luminescence/Fluorescence) ratio representing relative presynaptic ATP levels measured by the Syn-ATP sensor in mCherry positive varicosities of distal axons of DIV8 ctrl, KO, and KO+NAD^+^ neurons. Numbers (neurons imaged) and statistics: 78 ctrl, 77 KO, and 48 KO+NAD^+^ under basal conditions; 67 ctrl, 68 KO, and 48 KO+NAD^+^ under oligomycin-treated conditions. Ctrl and KO were from 5 independent experiments, and KO+NAD^+^ was from 2 independent experiments. Kruskal-Wallis test with Dunn’s multiple comparisons test was used. All above data represents mean ± SEM, ** p=0.004, ****p<0.0001. In the NAD^+^ treated KO group, the difference between basal and oligomycin treatment condition is marginal to significance, p= 0.0537.

NMNAT2 KO neurons exhibited a modest but significant decrease in sv-ATP compared to control neurons (Fig. 5C,D). sv-ATP can come from local glycolysis and/or mitochondrial ATP synthesis [82]. To check if it was contributed by glycolysis, we applied oligomycin, an F_1_-F_0_ ATP synthase inhibitor that blocks mitochondrial ATP production (Fig. 5A,B). Previous findings show that glycolytic ATP production is the primary ATP source for fast-transporting cargos [32, 81]. Here we only found a trend but not significant reduction of sv-ATP levels (p=0.3621) in control distal axons upon oligomycin treatment (Fig. 5C,D). However, in KO distal axons, oligomycin treatment significantly and strongly reduced sv-ATP levels (Fig. 5C,D), suggesting that in the absence of NMNAT2, mitochondria provide ATP in distal axons. These data strongly indicate that NMNAT2 is required to drive glycolysis on synaptic vesicles.

To test if the reduced sv-ATP in NMNAT2 KO axons is caused by NAD^+^ deficiency, neuronal cultures were supplemented with 1 mM NAD^+^ from DIV5 to DIV8 (Fig. 5A). Isotope labeling studies show that primary neuronal cultures can take up exogenous NAD^+^ [83, 84]. Here we found that NAD^+^ supplementation increased intracellular NAD^+^ levels transiently in neuronal cultures (Fig. 5-S2) and thus we refreshed NAD^+^ daily. NAD^+^ supplementation restored sv-ATP levels in KO distal axons to control levels in both basal and oligomycin treated conditions (Fig. 5C-D). These data suggest that NMNAT2 synthesizes the NAD^+^ to drive glycolysis on synaptic vesicles.

### NAD^+^ supplementation restores APP transport via glycolysis in the absence of NMNAT2

Next, we determined whether NAD^+^ supplementation can restore fast axonal transport in NMNAT2 KO distal axons. Compared to vehicle treatment, NAD^+^ supplementation significantly decreased the percentage of stationary/dynamic pause events, increased the percentage of anterograde and retrograde events, and restored anterograde and retrograde velocities of APP transport (Fig. 6A-C). On the other hand, NAD^+^ supplementation of control neurons had minimal impact on APP axonal transport (Fig. 6A-C) and NAD^+^ abundance (Fig. 5-S2).

**Fig 6.**
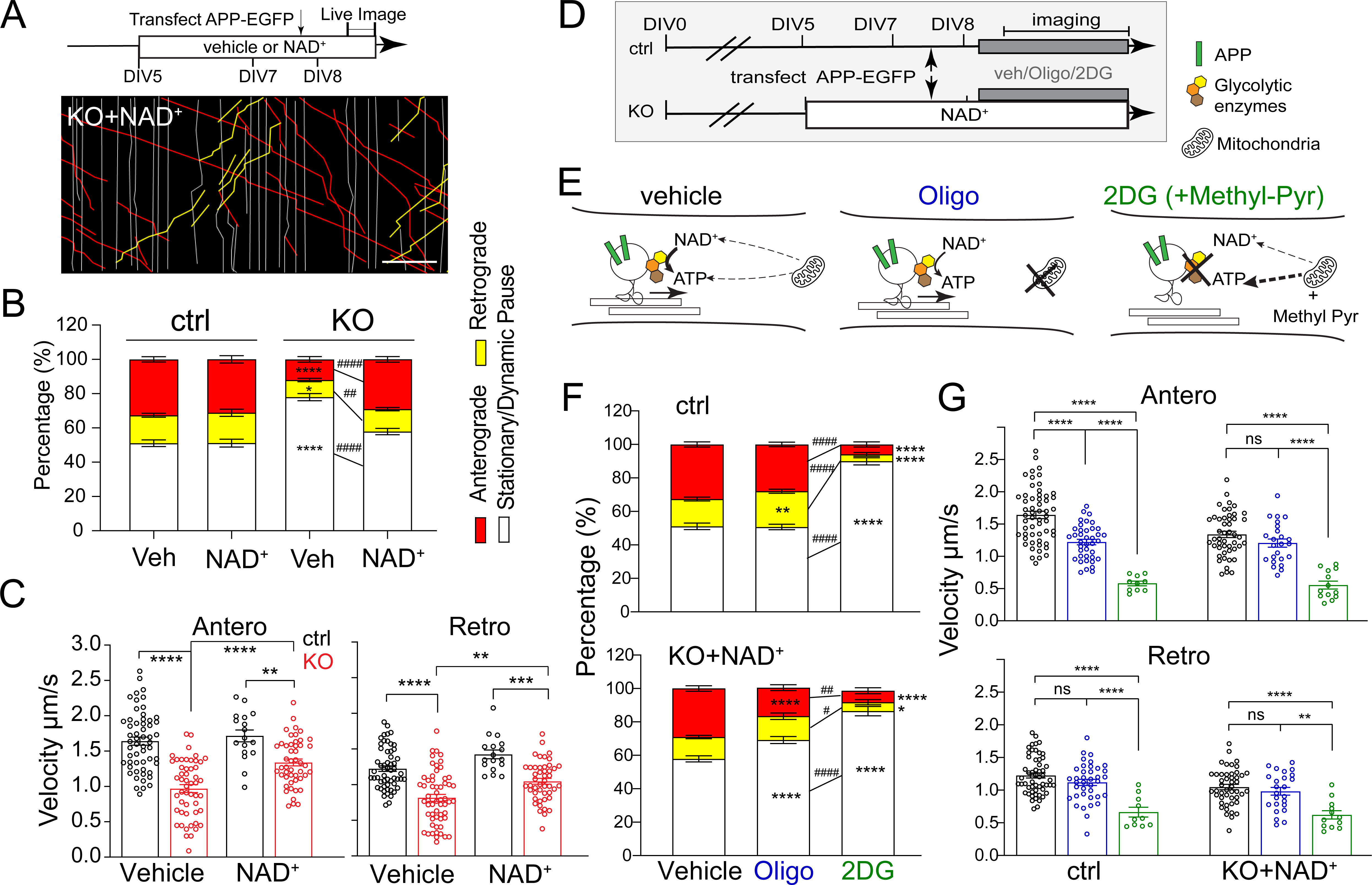
NAD supplementation depends on glycolysis to rescue APP transport in NMNAT2 KO neurons. **(A)** Representative kymographs from DIV8 ctrl or KO neurons supplemented with Veh (vehicle) or 1 mM NAD^+^ since DIV5. Scale bar, 20 µm. **(B)** Percentages of APP-EGFP vesicles moving anterogradely, retrogradely, or pausing in distal axons of DIV8 ctrl and KO neurons supplemented with Veh or NAD^+^. Numbers (neurons imaged) and statistics: 56 ctrl+Veh, 62 KO+Veh, 17 ctrl+NAD^+^,and 49 KO+NAD^+^, *p= 0.0232, ^##^p= 0.0018. Significant comparison to ctrl+Veh and KO+Veh groups was labeled with* and ^#,^ respectively. **(C)** Velocity of antero-and retrograde transport of APP-EGFP in distal axons of DIV8 ctrl and KO neurons supplemented with Veh or NAD^+^. Numbers (neurons imaged) and statistics: 56 ctrl+Veh, 51-58 KO+Veh, 17 ctrl+NAD^+^, and 47-48 KO+NAD^+^ for anterograde velocity, (anterograde) **p= 0.0041, (retrograde) **p= 0.0013, ***p= 0.0002. **(E)** Schematic representation of how glycolysis and mitochondrial respiration are affected by Oligo or 2DG(+Methyl-Pyr) treatment. **(F)** Percentages of APP-EGFP vesicles moving anterogradely, retrogradely or pausing in distal axons of DIV8 ctrl and KO+NAD^+^ neurons in the presence of vehicle, Oligo, or 2DG. Numbers (neurons imaged) and statistics: 56 ctrl+Veh, 39 ctrl+oligo, and 20 ctrl+2DG, **p=0.0059; 49 KO+NAD^+^+Veh, 22-25 KO+NAD^+^+Oligo,and 15-18 KO+NAD^+^+2DG, *p= 0.0291, ^#^p= 0.0252. Significant comparisons to Veh-treated and Oligo-treated groups were labeled with * and ^#,^ respectively. **(G)** Velocity of antero-and retrograde transport of APP-EGFP in distal axons of DIV8 ctrl and KO+NAD^+^ neurons in the presence of vehicle, Oligo, or 2DG. Numbers (neurons imaged) and statistics: 56 ctrl+Veh, 38-39 ctrl+oligo, 10 ctrl+2DG, 47-48 KO+NAD^+^, 24 KO+NAD^+^+Oligo, and 12-13 KO+NAD^+^+2DG, **p=0.0011. All above data represent mean ± SEM, two-way ANOVA with Tukey’s multiple comparisons test or Kruskal-Wallis test with Dunn’s multiple comparisons test were used, see details in supplemented excel, ****p<0.0001, ^####^p<0.0001.

We then tested whether glycolysis is required for NAD^+^ rescue of APP transport. We acutely suppressed glycolysis by transferring the neurons to a glucose-free medium containing the hexokinase inhibitor, 2-deoxyglucose (2DG), while adding methyl-pyruvate (Methyl-pyr) as an alternative substrate to support TCA cycle and OXPHO in mitochondria (Fig. 6D,E). In parallel, we tested if mitochondrial ATP production is required for NAD^+^ rescue of KO axonal transport by adding oligomycin in the presence of glucose to block OXPHO (Fig. 6D,E). Glycolysis inhibition significantly impaired APP transport in control axons, as revealed by a significant increase in stationary/dynamic pause events and decrease in transport velocity (Fig. 6F,G). In fact, the substantially reduced number of mobile APP-tagged vesicles in the glycolysis-inhibited cultures made it challenging to acquire sufficient vesicles for velocity measurements. In contrast, OXPHO inhibition had a relatively mild impact on APP transport in control axons, including a moderate reduction in anterograde velocity and a minor but significant increase in the percentage of retrograde events. In KO axons, glycolysis inhibition abolished the rescue of APP transport provided by exogenous NAD^+^, and significantly reduced the number of transport events and the movement velocities (Fig. 6F,G).

OXPHO inhibition also perturbed NAD^+^-mediated rescue of KO neurons, although not as robustly as did glycolysis inhibition, as shown by the increased percentage of stationary/dynamic pause events together and decreased percentage of anterograde events (Fig. 6F). However, the transport velocities in both directions were not affected (Fig. 6G). Additionally, we assessed APP accumulation by immunostaining and found that NAD^+^ supplementation from DIV8 to 14 significantly reduces APP accumulation in KO neurons. Furthermore, 48 hours of glycolysis inhibition together with methyl-pyruvate supplementation abolished this rescue (Fig. 6-S1A-C). These data demonstrate that NAD^+^-mediated rescue of APP axonal transport in NMNAT2 KO neurons depends mainly on glycolysis with a modest contribution from mitochondrial OXPHO.

### SARM1 depletion sustains APP transport and prevents axon degeneration in the absence of NMNAT2

Sterile alpha and TIR motif-containing protein 1 (SARM1) is a recently discovered NAD(P) glycol-hydrolase highly expressed in neurons [85, 86]. SARM1 senses the rise in nicotinamide mononucleotide (NMN)-to-NAD^+^ ratio caused by loss of NMNAT2 and responds to it by activating its own NAD^+^ hydrolase domain [87, 88]. SARM1 deletion in NMNAT2 KO mice prevents perinatal lethality, preserves healthy axons and synapses in the peripheral nervous system, and maintains motor function throughout adulthood [89, 90].

We therefore examined if SARM1 reduction could reverse axon phenotypes in NMNAT2 cKO brains. To this end, we crossed NMNAT2 cKO mice to SARM1 KO (S^null^/S^null^) mice to generate NMNAT2 cKO missing one copy of SARM1 (NMNAT2 cKO; S^null^/+), which we back-crossed to SARM1 KO mice to obtain NMNAT2 cKO missing both copies of SARM1 (NMNAT2 cKO; S^null^/S^null^). NMNAT2 cKO; S^null^/+ and NMNAT2 cKO; S^null^/S^null^ mice survived similarly to their littermate controls, in contrast to NMNAT2 cKO mice (data not shown). Normal brain morphology was observed in NMNAT2 cKO; S^null^/S^null^ mice (Fig. 7A). In contrast to NMNAT2 cKO; S^null^/S^null^ mice and control mice, NMNAT2 cKO; S^null^/+ mice still exhibited reduced body weights (Fig. 7-S1B). Gross examination of their brains found aberrant anatomy similar to NMNAT2 cKO brains (Fig. 7A), including atrophied hippocampi, enlarged ventricles, and thinned primary somatosensory cortex (Fig. 7B) and corpus callosum (Fig. 7C,D). APP accumulation was found in the corpus callosum, fimbria, and striatum in NMNAT2 cKO; S^null^/+ but not in NMNAT2 cKO; S^null^/S^null^ brains (Fig. 7E,F). APP accumulation was detectable at P5 in NMNAT2 cKO; S^null^/+ but not NMNAT2 cKO; S^null^/S^null^ brains, suggesting APP transport was already normalized by the complete absence of SARM1 at P5 (Fig. 7-S2). Taken together, these findings show that complete SARM1 loss prevents the impact of NMNAT2 loss in cortical axons.

**Fig 7.**
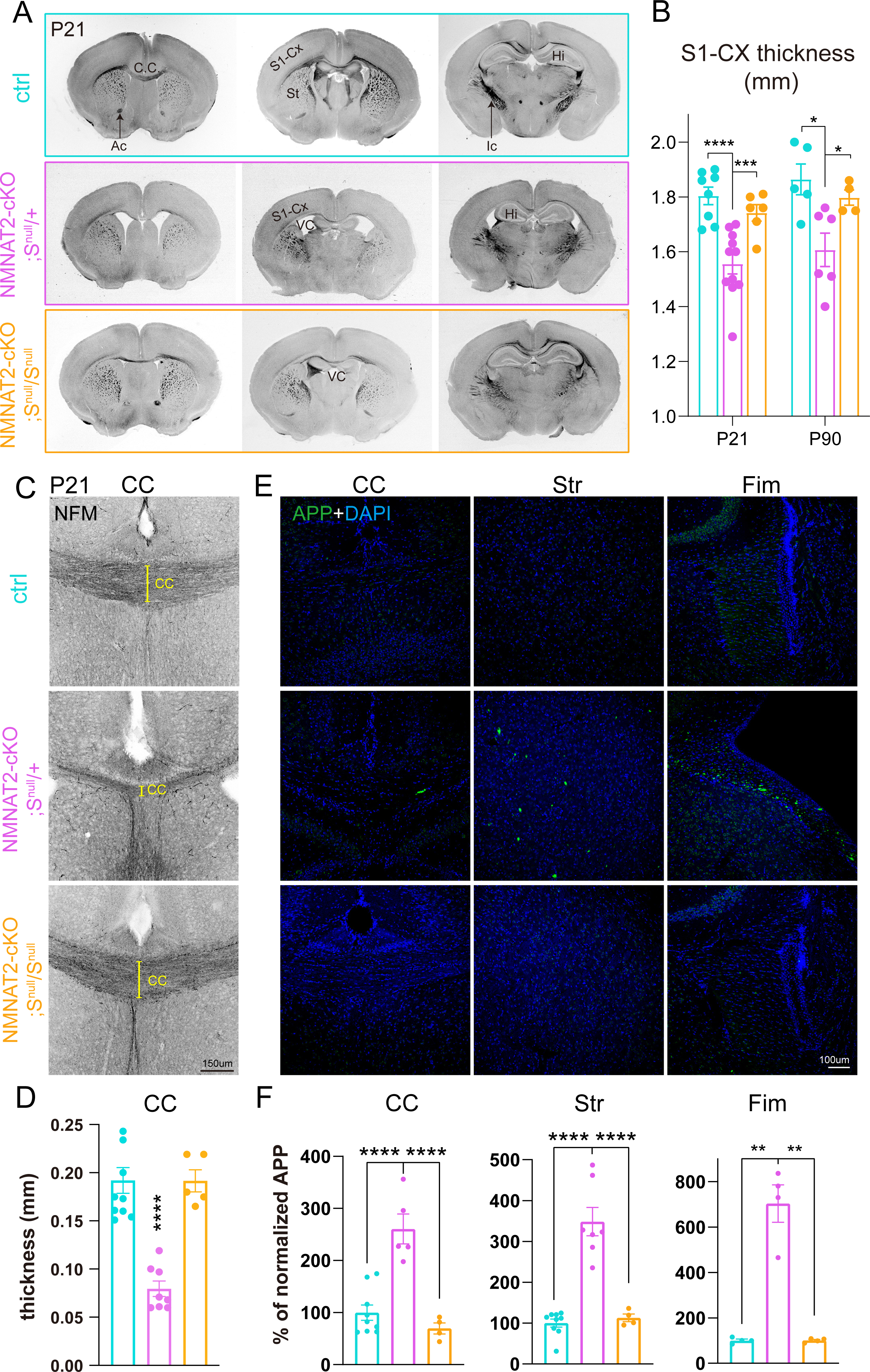
Deleting SARM1 prevents neurodegenerative phenotypes in NMNAT2 cKO brains. **(A)** Bright field images of coronal brain sections (rostral to caudal from left to right) from ctrl, NMNAT2 cKO; S^null^/+, and NMNAT2 cKO; S^null^/S^null^ mice. Enlarged ventricle, greatly reduced corpus callosum, and internal capsule were evident in NMNAT2 cKO; S^null^/+ but not in NMNAT2 cKO; S^null^/S^null^ brains. **(B)** Thickness of the primary somatosensory cortex in P16/21 and P90 mice. Number of mice: P16/21, 8 ctrl, 11 NMNAT2 cKO; S^null^/+, 6 NMNAT2 cKO; S^null^/S^null^; P90, 5 ctrl, 6 NMNAT2 cKO; S^null^/+, 4 NMNAT2 cKO; S^null^/S^null^. ***, p=0.0018, q=0.0009. (Ctrl vs. NMNAT2 cKO; S^null^/+) *, p= 0.0049, q= 0.0103. (NMNAT2 cKO; S^null^/+ vs. NMNAT2 cKO; S^null^/S^null^) *, p= 0.0341, q= 0.0358. **(C)** Representative NF-M (black) immunostaining images showing corpus callosum around the midline area in ctrl, NMNAT2 cKO; S^null^/+, and NMNAT2 cKO; S^null^/S^null^ rostral brain section at P21. **(D)** CC thickness quantification. Number of mice: P21, 10 ctrl, 8 NMNAT2 cKO; S^null^/+, 5 NMNAT2 cKO; S^null^/S^null^. **(E)** Immunohistology images showing APP accumulation in the corpus callosum, fimbria, and striatum of P21 NMNAT2 cKO; S^null^/+, but not in ctrl and NMNAT2 cKO; S^null^/S^null^ brains. (F) Quantification of the APP signal in the brain regions shown in **(E)**, normalized to the value in ctrl mice. Number of mice: 7-9 ctrl, 4-5 NMNAT2 cKO; S^null^/+, 4 NMNAT2 cKO; S^null^/S^null^. CC, corpus callosum; Str, striatum; Fim, Fimbria. (Fim Ctrl vs. NMNAT2 cKO; S^null^/+) **, p=0.0186, q= 0.0098. (NMNAT2 cKO; S^null^/+ vs. NMNAT2 cKO; S^null^/S^null^) **, p=0.0186, q=0.0098. All above data represent mean ± SEM. One-way ANOVA with two-stage linear step-up procedure of Benjamini, Krieger and Yekutieli was used, ****p<0.0001.

Next, we examined whether SARM1 loss protects NMNAT2 KO neurons from axonal degeneration by preventing axonal transport deficit. We used anti-sense oligonucleotides targeting sarm1 mRNA (SARM1-ASO) to knockdown SARM1 expression at desired time points, and a scrambled anti-sense oligonucleotide as the non-targeting control (ctrl-ASO). We evaluated the SARM1 knockdown efficacy of two SARM1-ASOs and one ctrl-ASO using qPCR and western blotting. Only ASO33, one of the two SARM1-ASOs, significantly decreased SARM1 protein abundance 4-7 days following ASO treatment (Fig. 8-S1). This ASO was therefore used for the rest of the experiments and referred to as SARM1-ASO. To test the impact of SARM1 knockdown on axonal transport, we applied ASOs from DIV1 and DIV5 on and examined the impact at DIV8. SARM1-ASO application starting at DIV1 significantly reduced SARM1 abundance by ∼70% and prevented APP transport deficits in NMNAT2 KO axons at DIV8 (Fig. 8A1-4). By contrast, SARM1-ASO treatment starting at DIV5 only reduced SARM1 abundance by ∼50% and failed to rescue axonal transport (Fig. 8B1-4). Surprisingly, by DIV12, APP transport was completely rescued in the distal axons of these neurons (Fig. 8C1-4), despite the impairment at DIV8. Consistent with axonal transport analysis, APP accumulation in KO neurons examined at DIV14 was rescued by SARM1-ASO treatment starting from DIV1 or DIV5 Consistent with axonal transport analysis, APP accumulation in KO neurons examined at DIV14 was rescued by SARM1-ASO treatment starting from DIV1 or DIV5 (Fig. 8-S2). Furthermore, the axon degeneration phenotype revealed by TUJ1 aggregates at DIV14 in ctrl-ASO-treated NMNAT2 KO axons was also reduced by SARM1-ASO treatment. Thus, blocking NAD^+^ degradation by SARM1 depletion protects axons during NMNAT2 loss in vivo and in vitro.

**Fig 8.**
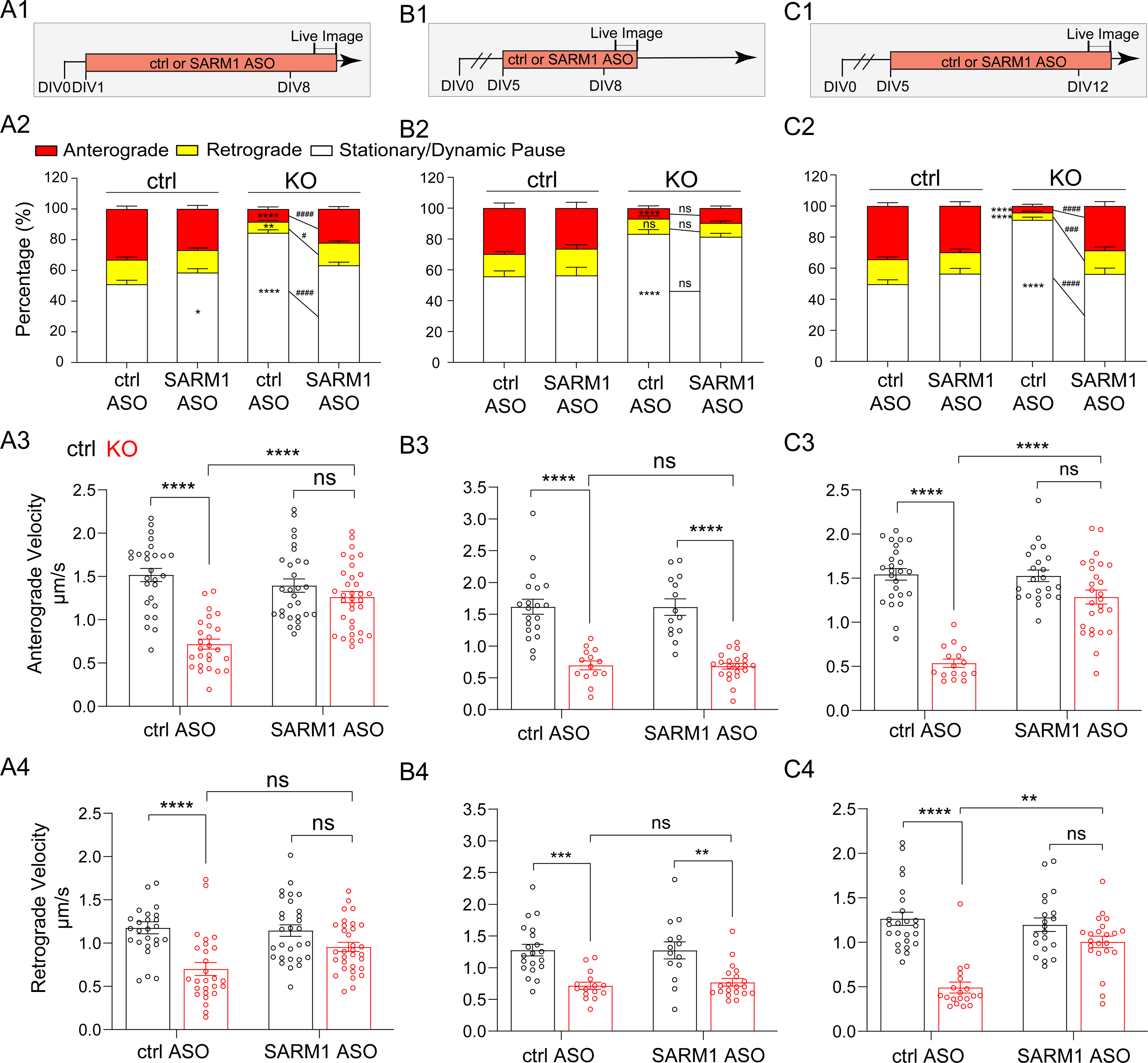
SARM1 knockdown restores APP axonal transport in NMNAT2 KO neurons. **(A1, B1, C1)** Experimental timeline. **(A2, B2, C2)** Percentages of APP vesicles moving anterogradely, retrogradely or staying stationary in distal axons of ctrl and KO neurons following the treatments indicated in **A1, B1**, and **C1**, respectively. Numbers of neurons imaged and statistics: **(A2)** 27 ctrl+ctrl ASO, 31 KO+ctrl ASO, 29 ctrl+SARM1 ASO, and 33 KO+SARM1 ASO, *p= 0.0286, ^#^p= 0.018, **p= 0.0069 . **(B2)** 20 ctrl+ctrl ASO, 22 KO+ctrl ASO, 14 ctrl+SARM1 ASO, and 26 KO+SARM1 ASO. **(C2)** 25 ctrl+ctrl ASO, 31 KO+ctrl ASO, 22 ctrl+SARM1 ASO, and 26 KO+SARM1 ASO, ^###^p=0.0006. Significant comparison to KO+ctrl ASO group was labeled with ^#^. **(A3-4, B3-4, C3-4)** Velocity of antero-and retrograde transport in distal axons of ctrl and KO neurons following the treatments indicated in **(A1, B1, C1)** respectively. **(A3-4)** 26-27 ctrl+ctrl ASO, 26-27 KO+ctrl ASO, 29 ctrl+SARM1 ASO, and 32-33 KO+SARM1 ASO. **(B3-4)** 20 ctrl+ctrl ASO, 14-16 KO+ctrl ASO, 14 ctrl+SARM1 ASO, and 20-22 KO+SARM1 ASO, (ctrl+SARM1 ASO vs KO+SARM1 ASO) **p=0.0049, (ctrl+ctrl ASO vs KO+ctrl ASO) **p=0.0022. **(C3-4)** 24-25 ctrl+ctrl ASO, 16-19 KO+ctrl ASO, 20-22 ctrl+SARM1 ASO, and 22-26 KO+SARM1 ASO, **p= 0.001. One-way or two-way ANOVA with Tukey’s multiple comparisons test and Kruskal-Wallis test with Dunn’s multiple comparisons test were used; see details in supplemented excel. All above data represent mean ± SEM, ****p<0.0001, ^####^p<0.0001. Numbers of independent culture experiments are listed in Sup. Table.

## Discussion

NMNAT2 has been identified as an AD therapeutic target and an axonal maintenance factor. Here we provide the first in vivo evidence that NMNAT2 is critical for the health of long-range cortical glutamatergic axons in mice. We show that NMNAT2 loss impairs glycolysis, disrupts fast axonal transport, and results in APP accumulation. NAD^+^ supplementation or reducing the levels of SARM1, an NAD^+^ hydrolase, effectively restores fast axonal transport and prevents the neurodegeneration normally observed in NMNAT2 cKO axons both in vitro and in vivo. In summary, our study suggests that NMNAT2 protects cortical neurons from axonal degeneration by ensuring that the energetic demands of distal axons are met. Our data also suggest that glucose hypometabolism in long-range axons is likely a major cause for axonopathy during NDAs. Our data indicate that therapies that prevent NMNAT2 or NAD loss during aging or NDAs could maintain axonal energetics to ensure axonal health.

### NMNAT2 contributes to fast axonal transport by maintaining the local NAD redox potential for efficient vesicular glycolysis

In this study, we show that NMNAT2 is required for maintaining vesicular glycolytic activity (Fig. 5) and the fast axonal transport of APP and SNAP25-containing vesicular cargos (Fig. 3) in distal axons of cortical neurons. This role in vesicular cargo transport is consistent with earlier observations. For instance, NMNAT2 is enriched in the synaptic vesicle and membrane, but not in mitochondrial fractions of cortical neurons [91]. In addition, palmitoylation of NMNAT2 enables it to associate with the membrane of Golgi-derived, fast-moving vesicular cargos [25]. Our time lapse imaging confirms that NMNAT2’s dynamic antero- and retrograde movement and transport velocities in axons are similar to those of fast transported cargos like APP- and SNAP25-containing vesicles (Fig. 3-S1).

Glycolysis is the major ATP source for fast axonal transport and is generated by a vesicular glycolytic complex [32, 81]. It has been estimated that one Kinesin requires ∼187.5 ATP molecules/s to move cargo at ∼1.5 μm/s [36], the average anterograde velocity of APP. Since each glycolysis reaction yields 2 ATPs, it must cycle ∼94 times per second near each cargo to power Kinesin, which is estimated to consume ∼188 molecules of NAD^+^. However, according to in vitro biochemical studies, NMNAT2 synthesizes at most ∼0.6 NAD^+^ per second [91]. Given that lactate dehydrogenases (LDH) catalyze NADH to NAD^+^, we hypothesize that additional NAD^+^ comes from LDH’s effort in recycling NADH to NAD^+^. Regardless, our data suggest that the NAD^+^ synthesizing activity of NMNAT2 and its proximity to the vesicular glycolytic complex ensure that enough ATP is produced to support motor proteins’ operation in fast axonal transport.

Intriguingly, before DIV8, NMNAT2 seems to be dispensable for axonal extension and axonal transport, which also requires glycolytic ATP supply [92]. We speculate that other mechanisms exist in the early developmental stage to ensure NAD homeostatic balance in the absence of NMNAT2, such as the newly discovered Wnk kinases [93]. However, by DIV8 in vitro and P16/21 in vivo, NMNAT2 becomes the mandatory source of NAD^+^, but only in distal axonal segments. In proximal axons, NMNAT2 is dispensable for NAD redox potential maintenance and fast axonal transport. Given that the NAD^+^ pool between the nucleus and cytoplasm are interchangeable [45], proximal axons may receive sufficient NAD^+^ from nuclear NMNAT1 due to their proximity to soma and thus maintain an adequate NAD redox potential to drive proximal axonal transport despite the absence of NMNAT2. Alternatively, OXPHO from somatic mitochondria could fuel axonal transport in proximal axons of NMNAT2 KO neurons. Consistent with this hypothesis, electron microscopy studies for cortical layers 2/3 of the primary visual cortex find that the proximal axons within 100 μm of the somata of pyramidal neurons tend to have higher mitochondrial coverage and volume than the distal axons do [94]. In distal axons, mitochondria tend to be smaller and less mobile [94–97]. We hypothesize that distal axon mitochondria are unable to compensate for the glycolysis deficits caused by loss of NMNAT2, which would explain why fast axonal transport in distal axons depends on NMNAT2.

Previous studies indicate that BDNF axonal transport is mainly fueled by glycolysis, and unaffected by ablating mitochondrial OXPHO [32]. Our data is largely consistent with this finding on glycolysis-dependent fast axonal transport. Nevertheless, we did observe, upon OXPHO inhibition, a mild but significant increase in the number of APP cargos undergoing retrograde transport, with a concomitant decrease in anterograde velocity. Using Drosophila neurodegenerative models, it has been shown that mitochondrial dysfunction stimulates a retrograde signaling response [98]. APP is partially localized on the endo-lysosomal vesicles involved in retrograde transport [99, 100]. Thus, the increase in retrograde APP-cargo percentages upon mitochondria inhibition may reflect enhanced retrograde signaling from the disrupted mitochondria in mouse glutamatergic cortical neurons as well.

Accumulation of APP, a pathological landmark for defective axonal transport [67, 101], was found in regions enriched in long-range axons of NMNAT2 cKO brains as early as P5, before obvious axonal loss. No APP accumulation was observed in the cortex, where proximal or short axonal arbors are enriched. Using cultured neurons, we demonstrated that the fast axonal transport of APP and SNAP25 is severely impaired in distal axonal segments of NMNAT2 KO neurons. These in vitro observations offer a mechanistic explanation for the APP accumulation in long-range axons observed in cKO brains. Together, our study provides strong evidence that NMNAT2 is required for fast axonal transport in distal axons of cortical glutamatergic neurons.

Cumulative evidence suggests that neuronal subtypes with long and complex axonal arbors are particularly vulnerable to external insults [102, 103]. As NMNAT2 cKO brains reach the age of P16-P21, not only does APP accumulation drastically increase, but severe axon degeneration occurs, along with a reduction in cortical thickness, hippocampal atrophy, ventricle enlargement, and motor behavioral deficits, recapitulating major hallmarks of neurodegeneration. These findings support the hypothesis that defective axonal transport drives axonopathy in the pre-symptomatic phase of neurodegeneration [54, 104], which may serve as the focal basis for the gradual development and spread of secondary neuronal damage [105, 106]. However, we do not exclude a possible role of NMNAT2 in maintaining neuronal health in the dendritic and somatic compartments [107, 108], whose dysfunction could also contribute to neurodegenerative phenotypes in NMNAT2 cKO brains.

### Mitochondrial OXPHO as a mediator of axonal degeneration in NMNAT2 KO cortical neurons

Our sv-ATP imaging studies show significantly reduced ATP levels in NMNAT2 KO distal axons when mitochondrial function is blocked (Fig. 5). By contrast, in control cortical axons, mitochondrial inhibition has a neglectable impact on sv-ATP levels. These observations suggest that NMNAT2 loss results in a shift towards mitochondrial OXPHO to maintain sv-ATP levels, however not enough to fuel fast axonal transport. The literature suggests two mechanisms that could mediate this shift in CNS neurons: (1) Ca^2+^ signaling-dependent regulation of mitochondrial calcium uniporters and the malate-aspartate shuttle [109–111]; (2) The O-GlcNAcylation-dependent post-translational modification of the mitochondrial proteome and mitochondrial motility [112, 113]. It remains unclear why this metabolic shift fails to protect NMNAT2 KO neurons from axonal degeneration: at DIV14, NMNAT2 KO axons exhibit prominent blebbing, a degenerative-like phenotype (Fig. 8-S2B). It has been suggested that the cumulative oxidative stress and depletion of TCA cycle substrates from hyperactive mitochondrial OXPHO could be detrimental to neuronal survival in PD and AD animal models [114–116]. This raises the possibility that the excessive mitochondrial activity that attempts to compensate for defective glycolysis in NMNAT2 KO axons contributes to axon degeneration.

We suspect SARM1 depletion not only mitigates the deleterious impact of excessive OXPHO, but also sustains OXPHO by preventing the decline of the mitochondrial NAD^+^ pool, thus offering excellent rescue of NMNAT2 KO neurons. Endogenous SARM1 widely distributes along neurites as small puncta [47, 117]. Upon mitochondrial stress or overexpression, SARM1 localizes onto the mitochondrial outer membrane and interacts with mitophagy machineries [85, 86, 118, 119]. SARM1 is a NAD^+^ consuming enzyme and its NAD^+^ hydrolase activity can be activated by the rise of nicotinamide mononucleotide to NAD^+^ ratio [87], or by JNK-mediated phosphorylation [120].

It has been shown that SARM1 in an activated state drastically consumes NAD^+^ and impairs mitochondrial OXPHO [120]. In an axotomy model, mitochondrial motility is preserved when SARM1 is genetically deleted [121]. In a Charcot-Marie-Tooth neuropathy model with mitochondrial abnormality, SARM1 deletion preserved mitochondrial morphology and motility, and protected axons and synapses from degeneration [122]. Here we find that SARM1 loss prevents several phenotypes caused by NMNAT2 loss, including the impaired fast axonal transport and axonal morphology. Currently, it is still unknown whether SARM1 removal in NMNAT2 KO axons prevents the oxidative stress caused by mitochondria hyperactivity. Further investigations should examine the impact of deleting SARM1 from NMNAT2 KO axons on mitochondrial metabolic capacity, dynamics, NAD^+^-NADH shuttling between mitochondrial matrix and cytoplasm, and mitochondrial quality control.

### NMNAT2-dependent vesicular glycolysis as a potential target to protect white matter in neurodegenerative diseases

Positron emission tomography (FDG-PET) studies have shown that glucose metabolism changes during aging/NDAs in specific brain regions [123–128]. In particular, glucose hypometabolism is often detected in AD susceptible brain regions, and strongly predicts the incidence of mild-cognitive impairment (MCI) in later life [129–133]. The majority of AD-susceptible regions for cognitive decline identified are located in the default mode network (DMN), the topological central hub of the brain connectome [134–136]. Anatomically, these hub areas are densely inter-connected by long-range axonal tracts [137] that often show pathological changes since the early stages of disease [138]. Metabolically, these connectome hubs express higher levels of glycolysis genes compared to the non-hub areas [139], and exhibit reduced aerobic glycolysis upon normal aging and AD [140, 141], which positively correlates with white matter deterioration [142]. Notably, glucose hypometabolism is observed directly in white matter tracts of AD brains [4].

Here we show that NMNAT2 is critical for vesicular glycolysis to fuel axonal transport in distal axons. Previously, we have shown that NMNAT2 levels are significantly reduced in the prefrontal cortex of AD brains [21], one of the DMN hubs. Data from our biochemical assays reveal ∼50% reduction of NAD^+^ and NADH levels in NMNAT2 KO neurons (Fig. 4A), indicating that NMNAT2 is a major NAD^+^ and NADH provider in cortical neurons. Synthesizing these observations, we hypothesize that reduced NMNAT2 in AD brains contributes, at least in part, to the reduction in glucose metabolism observed in AD white matter.

NMNAT2 reduction has also been observed in PD and HD brains [21–23]. PD patients show hypometabolism in the premotor and parieto-occipital cortex that correlates with motor dysfunction [128], while HD patients show progressive glucose hypometabolism in the frontal lobe, temporal lobe, and striatum, accompanied by white matter volume reduction [125]. Supplementing NAD^+^ and its precursors provides remarkable amelioration of degenerative phenotypes in transgenic AD and ALS mouse models [143–145]. Similarly, increasing NMNAT2 expression broadly provides neuroprotection across mouse models of tauopathy [21, 146], familiar AD [147], and glaucoma [148]. Coincidently, upregulating glycolysis exerts neuroprotective effect in PD synucleinopathy and ALS models [149, 150]. Our study highlights the role of NMNAT2 in supporting glycolysis in long-range projecting axons of cortical glutamatergic neurons. Extrapolating from our findings, NMNAT2 could serve as a putative therapeutic target to boost neuronal glycolysis in order to antagonize the structural connectome breakdown occurring in many neurodegenerative disorders.

## Supporting information

Data and statistics

---

**Fig 1-S1.**
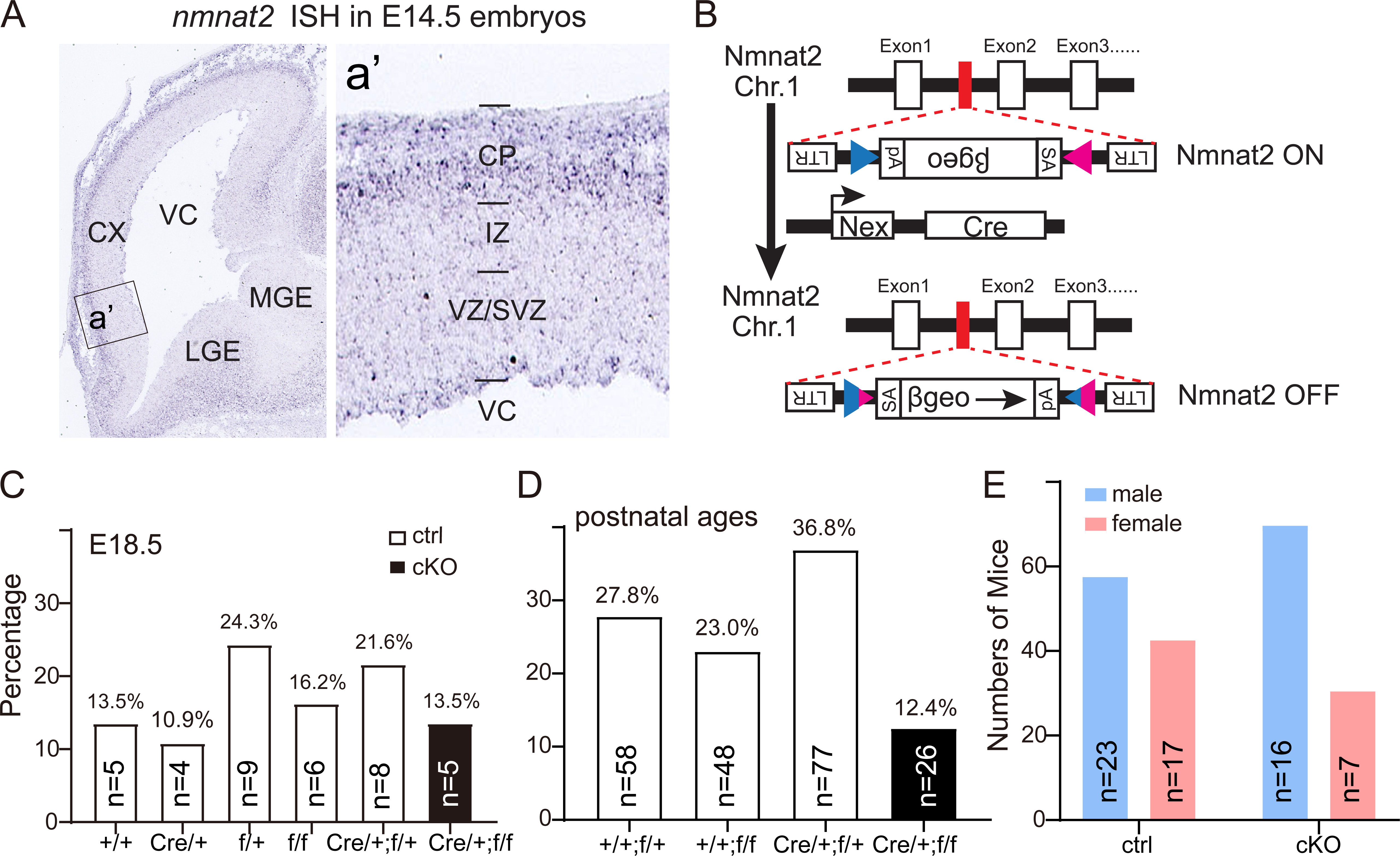
NMNAT2 expression and cKO mice survival rate. (**A**) In situ hybridization reveals nmnat2 mRNAs (blue signal) in the E14.5 brain, mainly in the cortical plate (CP) and in the lateral ganglionic eminence (LGE) and medial ganglionic eminence (MGE). (**B**) Illustration of Nmnat2 conditional KO (cKO) mice generation as described previously [37]. The blue and magenta triangles indicate loxP sites. (C, D) Summary of embryonic and postnatal pups genotyping. For embryonic samples, the strategy was to breed Nex-Cre/+;NMNAT2^f/+^ to with NMNAT2^f/+^ mice. The genotyping results conform to the expectations of Mendelian inheritance. The breeding strategy for postnatal pups was to cross Nex-Cre/+;NMNAT2^f/+^ with NMNAT2^f/f^ mice. The genotyping result showed a smaller than expected number of mice with the Nex-Cre/+;NMNAT2^f/f^ (cKO) genotype, suggesting a reduced survival rate for cKO mice after birth. (**E**) Overall numbers of postnatal male and female ctrl and cKO mice. Abbreviations: Cre, Nex-Cre; f/f, NMNAT2^f/f^; IZ, intermediate zone; Str, striatum; SVZ, subventricular zone; VC, ventricle; VZ, ventricular.

**Fig 2-S1.**
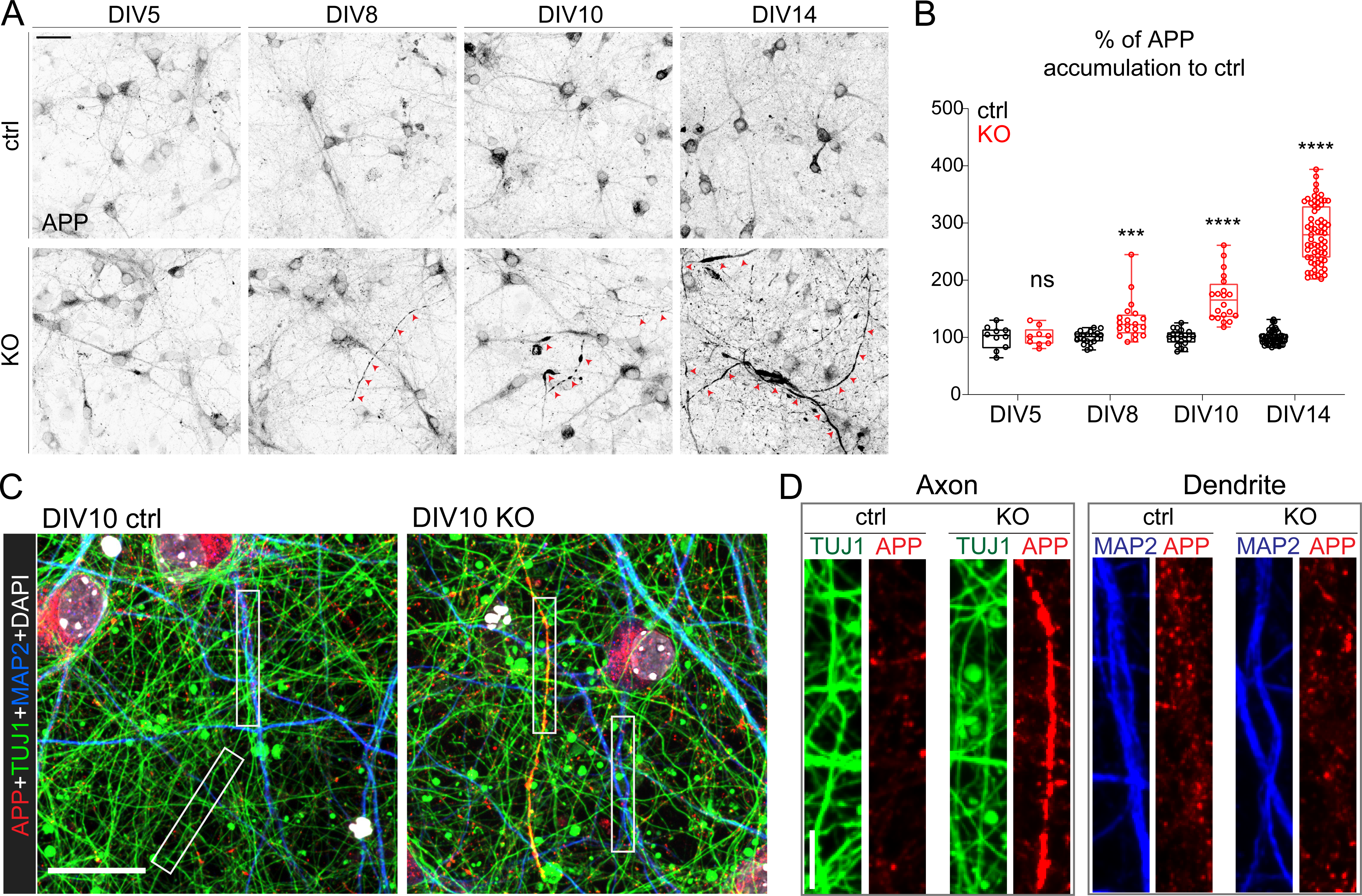
Loss of NMNAT2 triggers APP accumulation in the axon of cultured cortical neurons. (**A**) Representative confocal image of APP immunocytochemical staining in ctrl and KO neuronal culture at DIV5, 8, 10, and 14. Scale bar 30 μm. Red arrows highlight APP accumulation. (**B**) Area of accumulated APP in cKO and ctrl mice, normalized to the average ctrl value of the corresponding DIV. Image numbers used for analysis: DIV5: 10 ctrl, 9 KO; DIV8 & DIV10: 20 ctrl, 20 KO; DIV14: 50 ctrl, 67 KO. DIV5-10 were from 2 independent culture experiments and DIV14 was from > 3 independent experiments. Multiple Mann-Whitney test with Holm-Šídák post-hoc adjustment, KO was compared to ctrl of the same DIV, ***p=0.000202, ****p<0.0001. (**C**) Representative confocal images of APP, TUJ1 (βIII-tubulin), MAP2, and DAPI staining in DIV10 ctrl and KO neurons. Scale bar 20 μm. (**D**) Zoom-in view of axon and dendrite area highlighted by the white box in (**C**). Scale bar 5 μm.

**Fig 2-S2.**
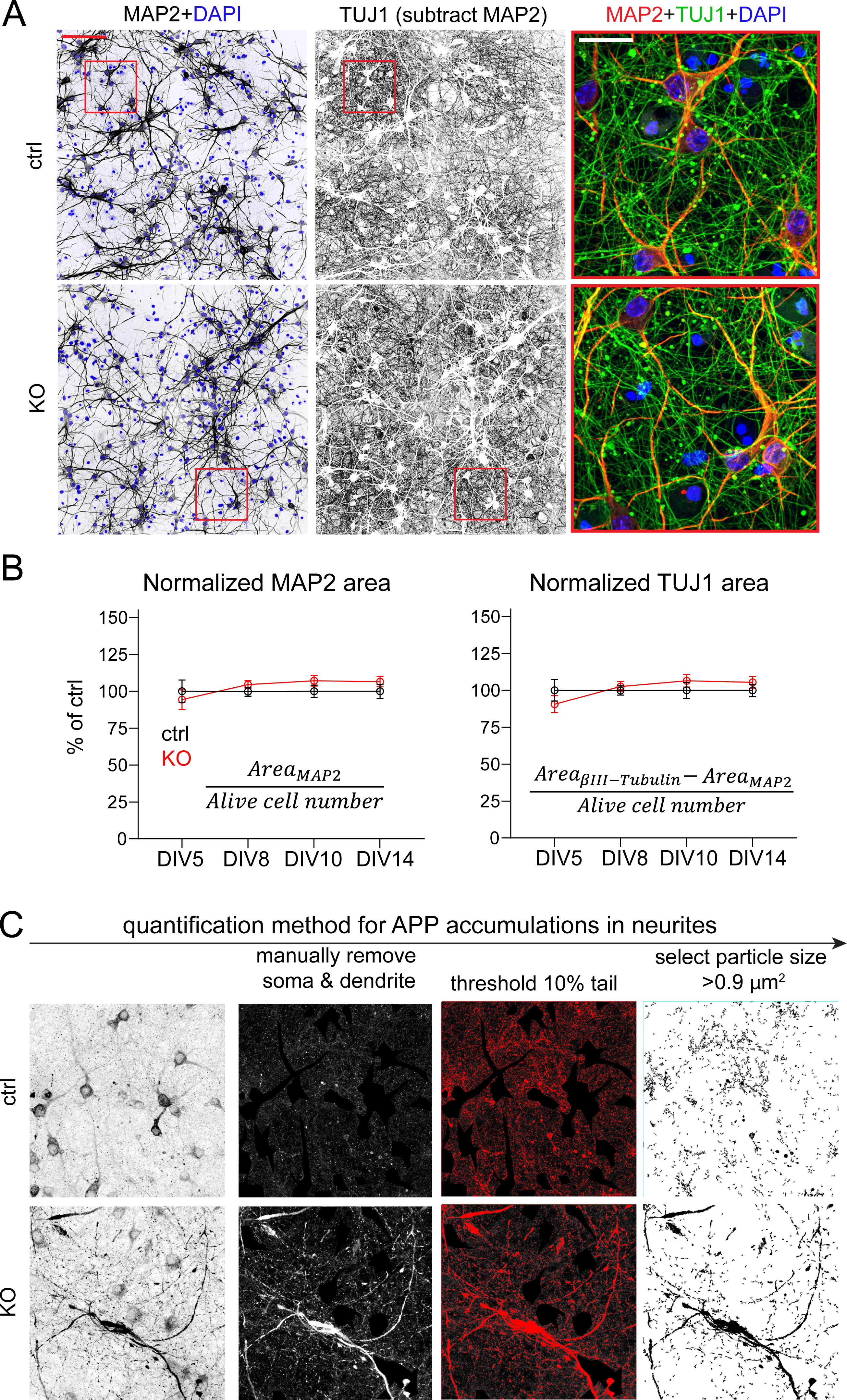
Gross neurite covering the area in vitro is not affected by Nmnat2 loss. (**A**) Representative confocal images of MAP2, TUJ1, and DAPI staining in ctrl and KO neuronal culture at DIV10. MAP2 positive area corresponding to soma and dendrites was subtracted in TUJ1 staining images to determine area corresponding to axon regions. 100 μm scale bar in left and middle panels; 25 μm scale bar in zoom-in panels. (**B**) Quantification of the somatic and dendritic area labeled by MAP2 and the axonal area labeled by TUJ1 (MAP2 area subtracted) in ctrl and KO neuronal culture at DIV5, 8, 10, and 14, normalized to the average ctrl values at the same DIV. Image numbers from 1-3 independent experiments: DIV5: 10 ctrl, 10 KO; DIV8: 23 ctrl, 16 KO; DIV10: 11 ctrl, 13 KO; DIV14: 5 ctrl, 6 KO. Multiple Mann-Whitney tests with Holm-Šídák correction for comparisons were used for MAP2; two-way ANOVA with Šídák’s multiple comparisons test was used for TUJ1. Data represent mean ± SEM. (**C**) The procedure of semi-automated quantification of accumulated APP by ImageJ.

**Fig 2-S3.**
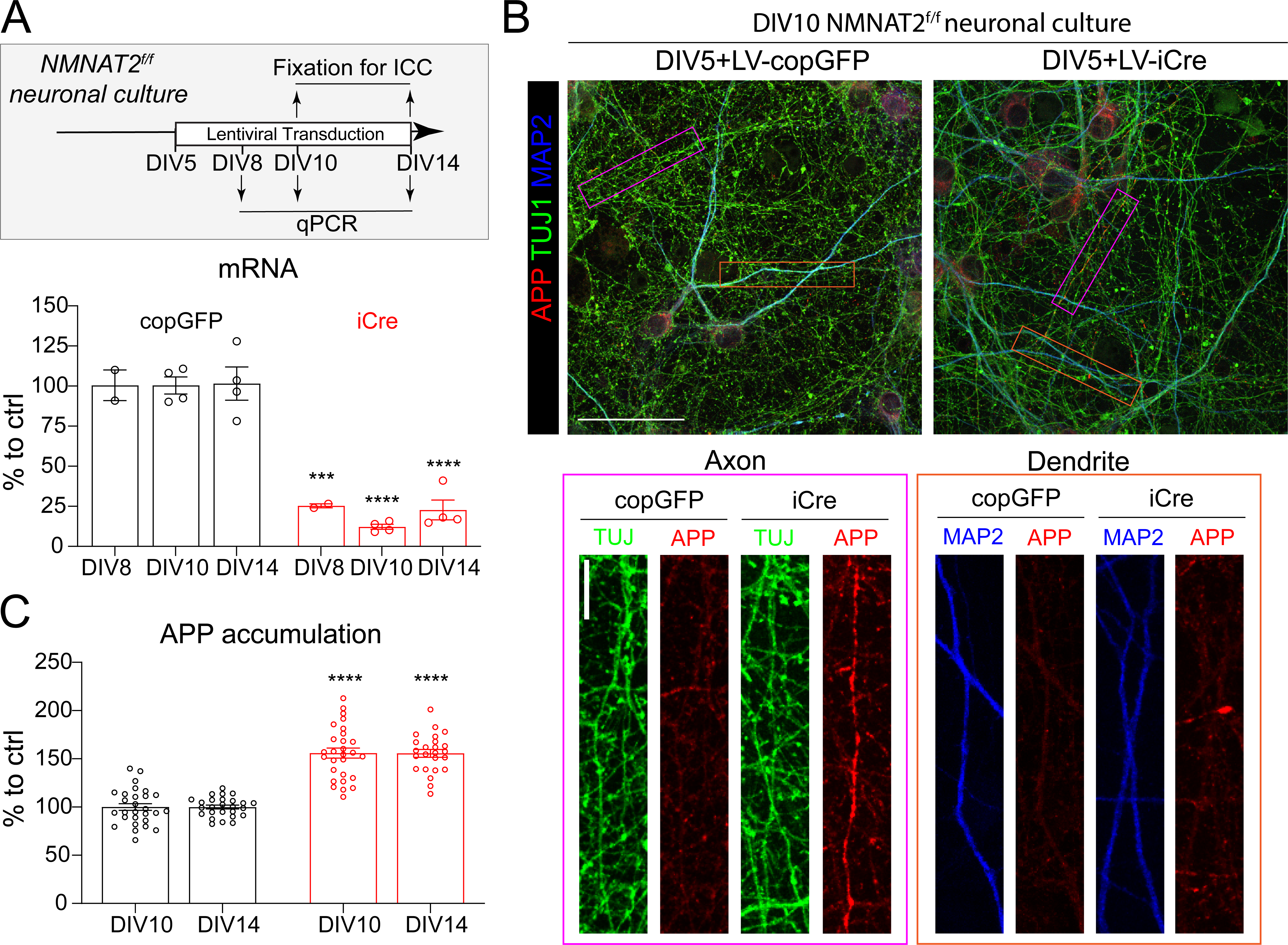
Loss of NMNAT2 after neurite outgrowth in vitro triggers axonal APP accumulation. (**A**) The experimental timeline in NMNAT2^f/f^ cortical neuronal culture. The NMNAT2 mRNA level was normalized to the LV-copGFP transduced ctrl group of the corresponding DIV and expressed as percentage. Numbers of culture wells from 1-2 independent batches of culture: DIV8: 2 copGFP, 2 iCre; DIV10 & 14: 4 copGFP, 4 iCre; (**B**) Representative confocal images of APP, TUJ1 and MAP2 staining in DIV10 NMNAT2^f/f^ neurons that received lentivirus starting at DIV5, 30 μm scale bar. Zoom-in view of axon and dendrite area highlighted by magenta and orange box respectively in the upper panel, 10 μm scale bar. (**C**) The area of accumulated APP was normalized to the LV-copGFP transduced neurons of the corresponding DIV as percentage. Number of images acquired from 2-3 independent experiments: DIV10: 28 copGFP, 28 iCre; DIV14: 26 copGFP, 24 iCre. Two-way ANOVA with Šídák’s multiple comparisons test, iCre was compared to copGFP of the same DIV, ****p<0.0001.

**Fig 3-S1.**
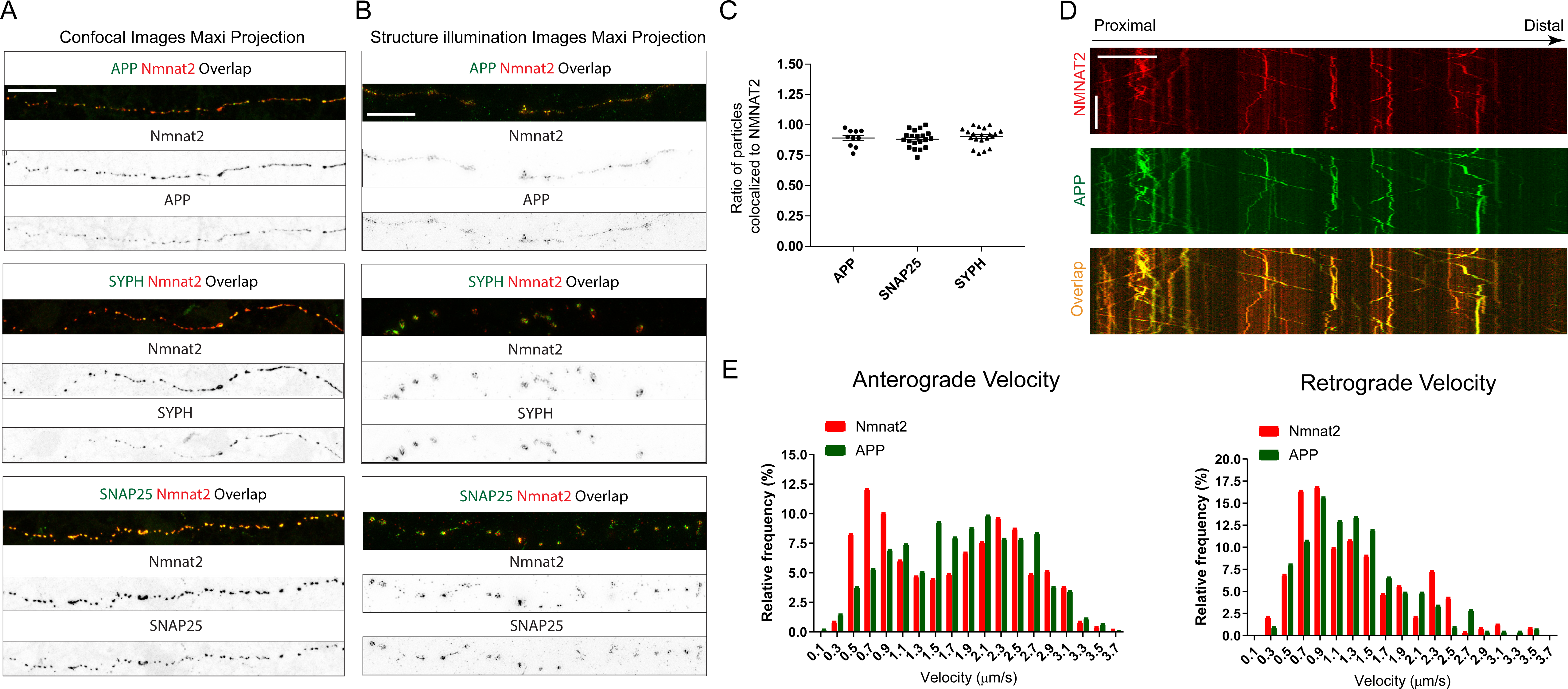
NMNAT2 colocalize and co-migrate with vesicular cargos. (**A**) Representative confocal images with maximum intensity Z stack projection showing that NMNAT2-mCherry colocalizes with APP-EGFP, SYPH-EGFP and SNAP25-EGFP in axons of DIV8 WT cortical neurons (Scale bar, 20 μm). (**B**) Representative structure illumination images with maximum intensity Z stack projection showing NMNAT2-mCherry colocalizes with APP-EGFP, SYPH-EGFP and SNAP25-EGFP in axons of DIV8 WT cortical neurons (Scale bar, 5 μm). (**C**) The ratio of the particles of APP-EGFP, SYPH-EGFP, and SNAP25-EGFP colocalized to NMNAT2-mCherry in (**A**). Number of neurons, APP (N=10), SNAP25 (N=20), and SYPH (N=20). (**D**) Representative kymograph showing NMNAT2-mCherry comigrates with APP-EGFP in axon of DIV8 WT neurons (Scale bar 20 μm in length, 30 s in time, 1.7 frame/second). (**E**) Frequency distribution of anterograde and retrograde velocity of NMNAT2-mCherry and APP-EGFP in DIV8 distal axons. (NMNAT2 N=445, APP N=861, N represents the cargo trajectory traced in all kymographs).

**Fig 3-S2.**
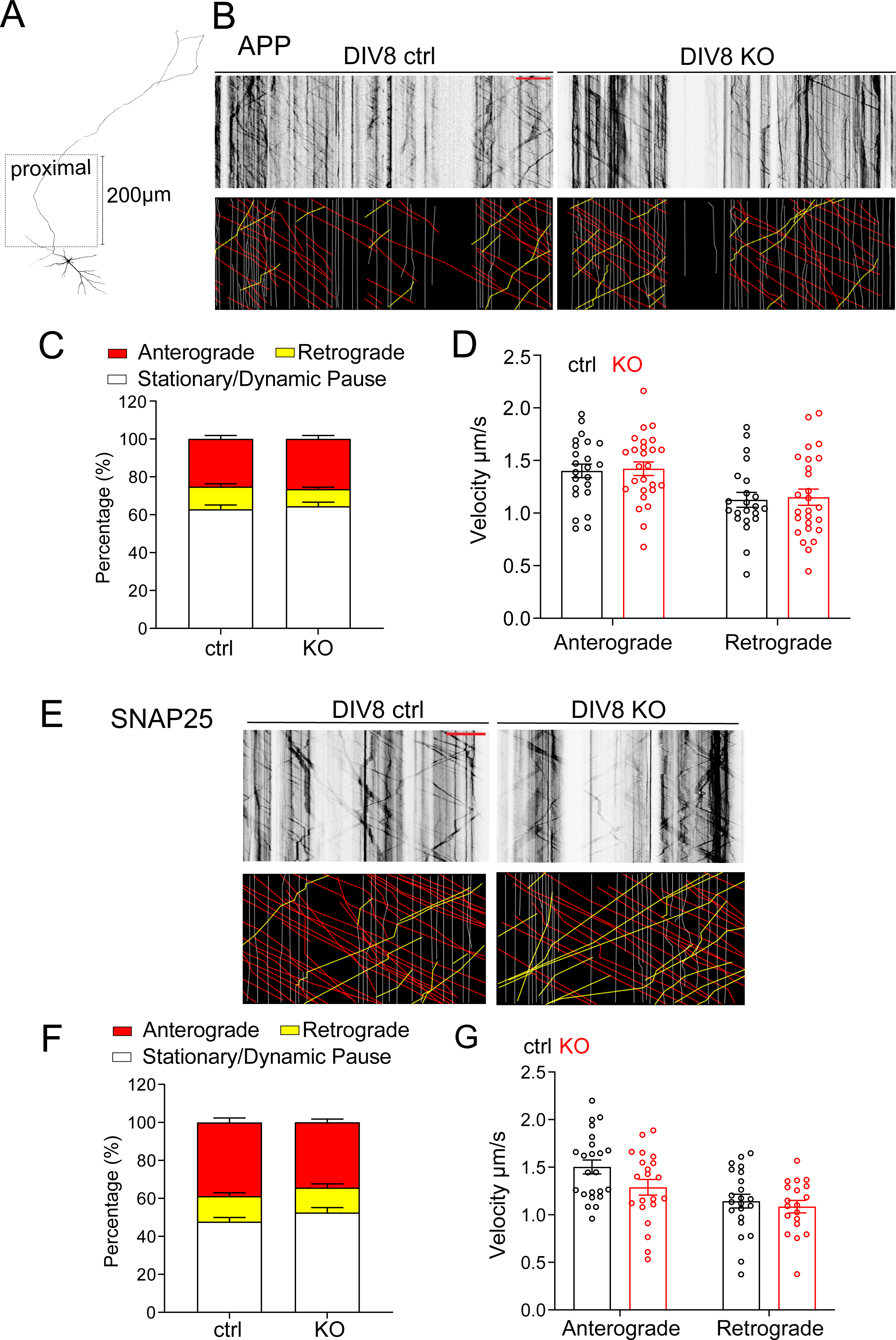
Vesicular cargo transport in the proximal axon is less affected by NMNAT2 loss at DIV8. (**A**) Diagram indicating the proximal axon region (< 200 µm from the soma). (**B**) Representative kymographs for APP-EGFP from DIV8 ctrl and KO proximal axons. Scale bar, 20 µm. (**C**) Distribution of APP-EGFP vesicles in anterograde motion, retrograde motion, or in a stationary/dynamic pause from 2 independent culture experiments. Numbers (neurons imaged) and statistics: 23 ctrl, 27 KO; compared with unpaired t-test. (**D**) Velocity of antero-and retrograde transport of APP-EGFP from 2 independent culture experiments. Numbers (neurons imaged) and statistics: anterograde, 23 ctrl, 26 KO; retrograde, 22 ctrl, 26 KO. Two-way ANOVA with Šídák’s multiple comparisons. (**E**) Representative kymographs for SNAP25-EGFP from DIV8 ctrl and KO proximal axons. Scale bar, 20 µm. (**F**) Distribution of APP-EGFP vesicles in anterograde motion, retrograde motion, or in a stationary/dynamic pause from 3 independent culture experiments. Numbers (neurons imaged) and statistics: 23 ctrl, 21 KO; anterograde and retrograde categories were analyzed by Mann-Whitney test, and stationary/dynamic pause category was analyzed by unpaired t-test. (**G**) Velocity of antero- and retrograde transport of SNAP25-EGFP from 3 independent culture experiments. Numbers (neurons imaged) and statistics: anterograde, 23 ctrl, 21 KO; retrograde, 22 ctrl, 20 KO. Two-way ANOVA with Šídák’s multiple comparisons. Data represent mean ± SEM.

**Fig 3-S3.**
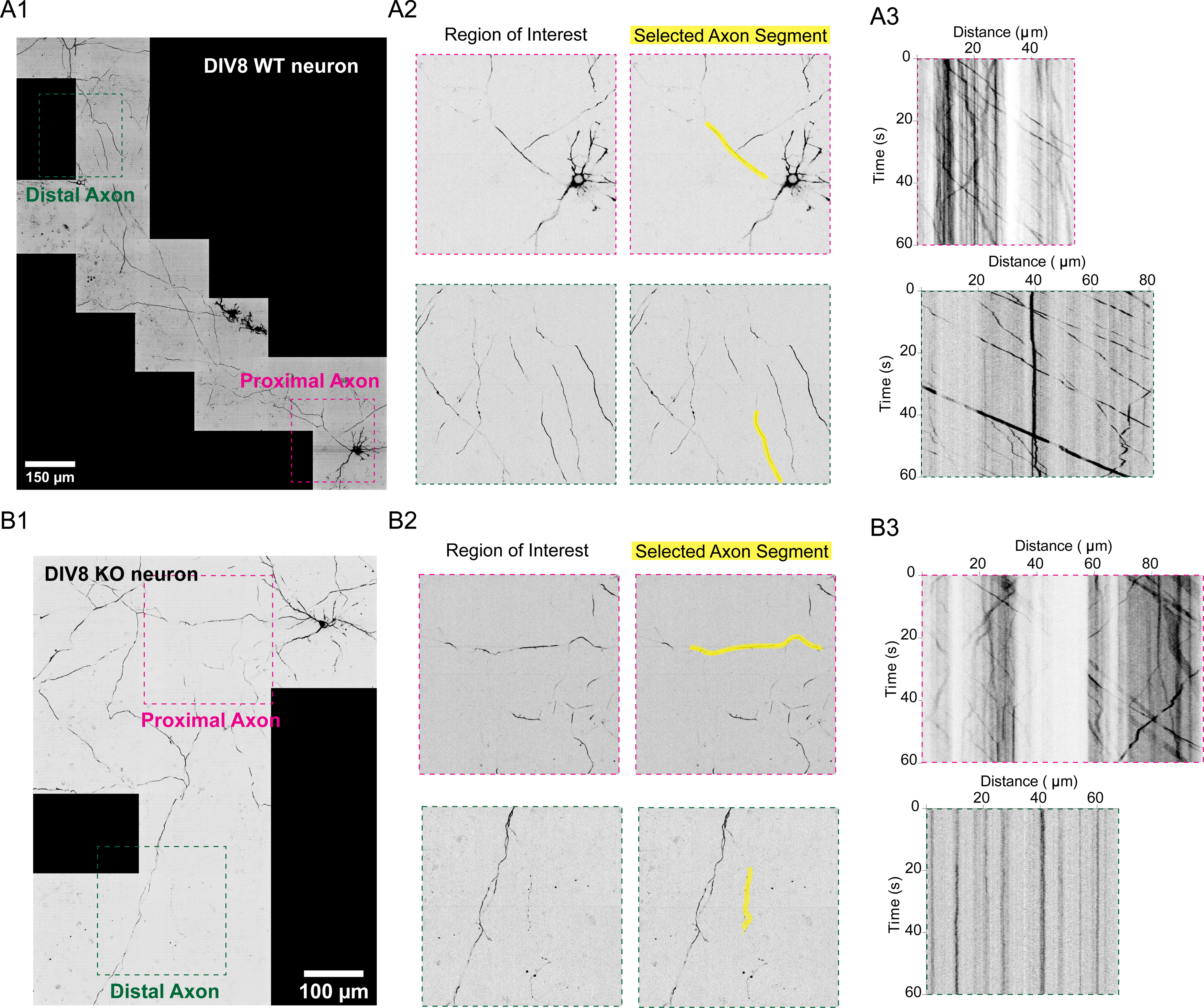
Examples of axonal morphology overview. (A1, B1) Stitched confocal image of DIV8 ctrl or KO neuron expressing SNAP25-EGFP. (A2, B2) Zoom-in of the regions of interest identified as proximal and distal axons as highlighted in magenta and dark green boxes in (A1, B1). (A3) Kymograph generated from the axon segment selected and highlighted in yellow in (A2, B2).

**Fig 3-S4.**
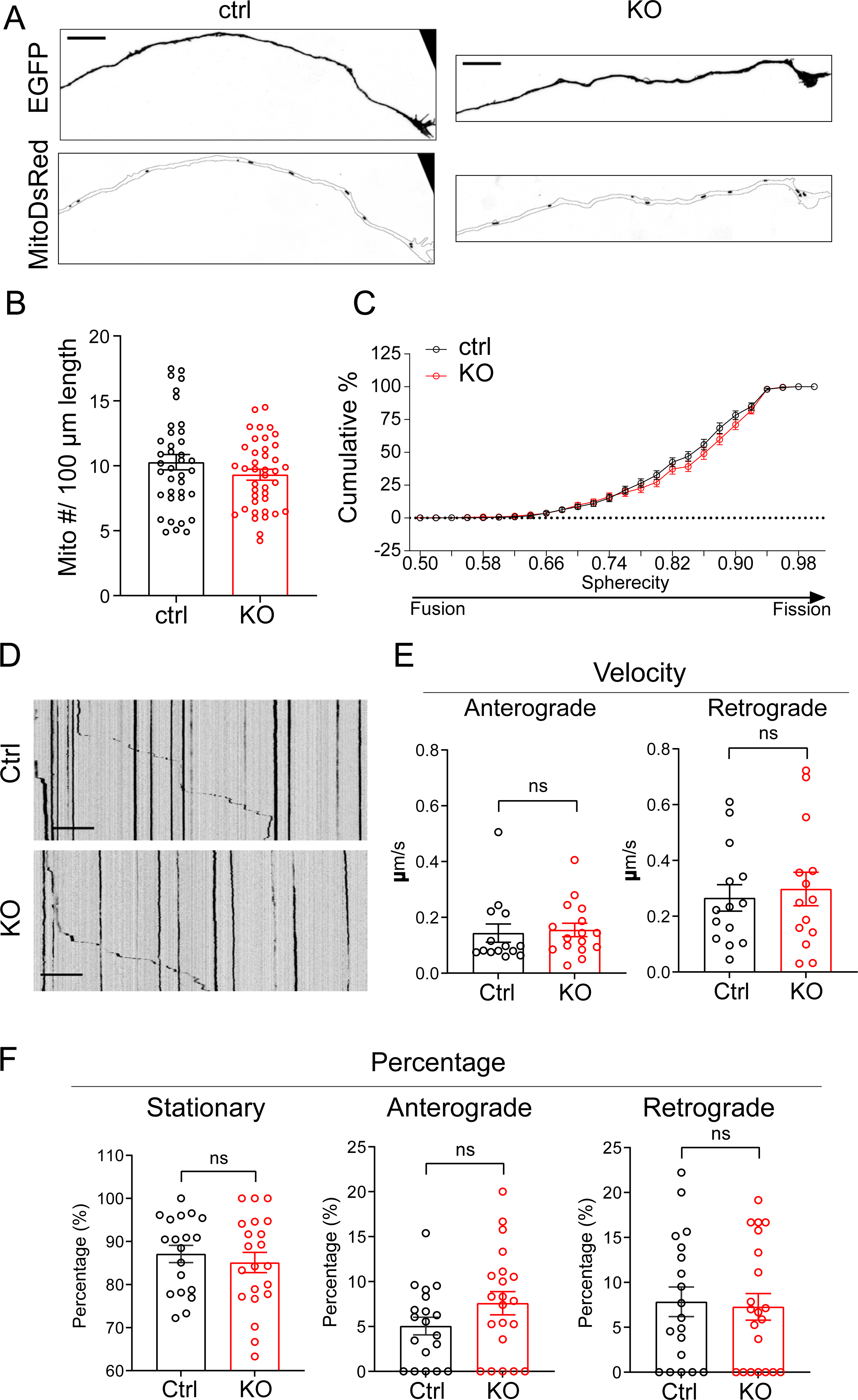
Mitochondrial density, morphology, and mobility in distal axons are unaffected by NMNAT2 loss at DIV8. (**A**) Representative images of EGFP and Mito-DsRed in the distal axon of DIV8 ctrl and KO neurons. (Scale bar 10 μm). (**B**) Mitochondria density (# of mitochondria per 100 μm) in distal axons of DIV8 ctrl and KO neurons from 3 independent experiments. Numbers (neurons imaged) and statistics: 38 ctrl, 40 KO; unpaired t-test was used. (**C**) The cumulative percentage distribution of mitochondria morphology indicated by sphericity in DIV8 ctrl and KO distal axons from 3 independent experiments. Numbers (neurons imaged) and statistics: 38 ctrl, 40 KO; Log(Y/(1-Y)) transformation for the independent variable and linear mixed model with random intercept and the slope was used; see descriptions in Star Methods. (**D**) Representative kymographs show the mobility of MitoDsRed in distal axons of DIV8 ctrl and KO neurons. (Scale bar, 20 µm; total recording time, 15min). (**E**) The velocity of anterogradely and retrogradely moving MitoDsRed in distal axons of DIV8 ctrl and KO neurons from 2 independent batches of culture. (Anterograde: ctrl N=14, KO N=16, Mann-Whitney test was used; Retrograde: ctrl N=14, KO N=14, unpaired t test was used). (**F**) Percentage of MitoDsRed that is anterogradely moving, retrogradely moving, or in stationary/dynamic pause in distal axons of DIV8 ctrl and KO neurons from 2 independent batches of culture (ctrl N=19, KO N=21). For analyzing anterograde and stationary/dynamic pause, an unpaired t-test was used; for retrograde, the Mann-Whitney test was used. N represents number of neurons imaged. All above data represents mean ± SEM.

**Fig 5-S1.**
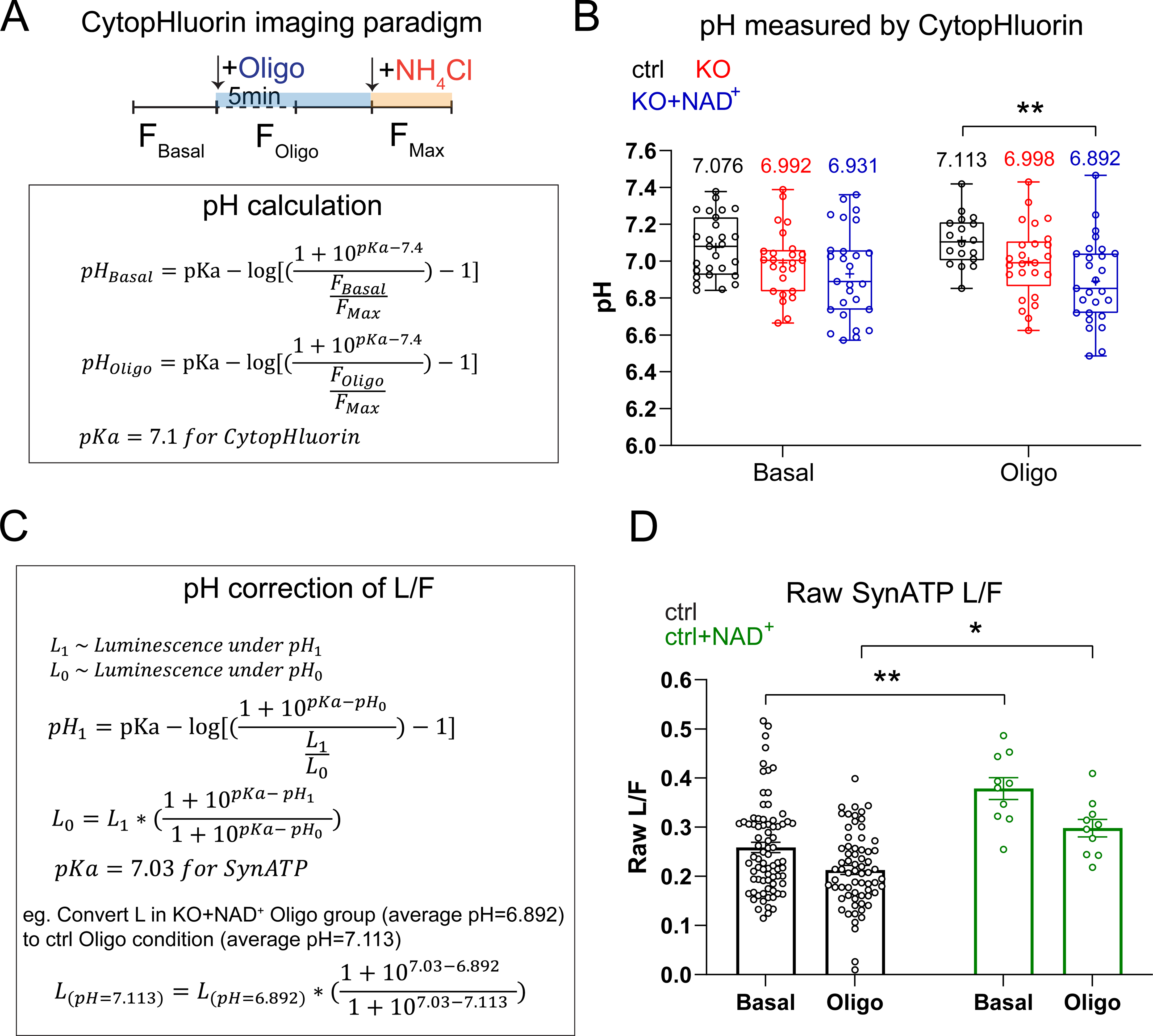
pH measurement and pH correction for Syn-ATP. (**A**) Cyto-pHluorin imaging timeline and equation used for cytosolic pH determination in presynaptic varicosities. (**B**) pH measured by Cyto-pHluorin in various conditions. The mean value in each group is labeled above each box and whisker. Numbers (neurons imaged) and statistics: 27 ctrl, 25 KO, and 27 KO+NAD^+^ under basal conditions; 18 ctrl, 25 KO, and 27 KO+NAD^+^ under oligomycin-treated conditions. All groups were from a single experiment. Two-way ANOVA with Tukey’s multiple comparisons test, **p= 0.005. (**C**) Equations used for pH correction of L/F measurements from Syn-ATP. (**D**) Raw L/F measurements in distal axon varicosities of DIV8 ctrl neurons with or without NAD^+^ supplementation. Numbers (neurons imaged) and statistics: 78 ctrl and 10 ctrl+NAD^+^ under basal conditions; 67 ctrl and 10 ctrl+NAD^+^ under oligomycin-treated conditions. Ctrl was from 5 experiments, and ctrl+NAD^+^ was from one experiment. Kruskal-Wallis test with Dunn’s multiple comparisons test, *p= 0.0171, **p= 0.0018. All above data represent mean ± SEM.

**Fig 5-S2.**
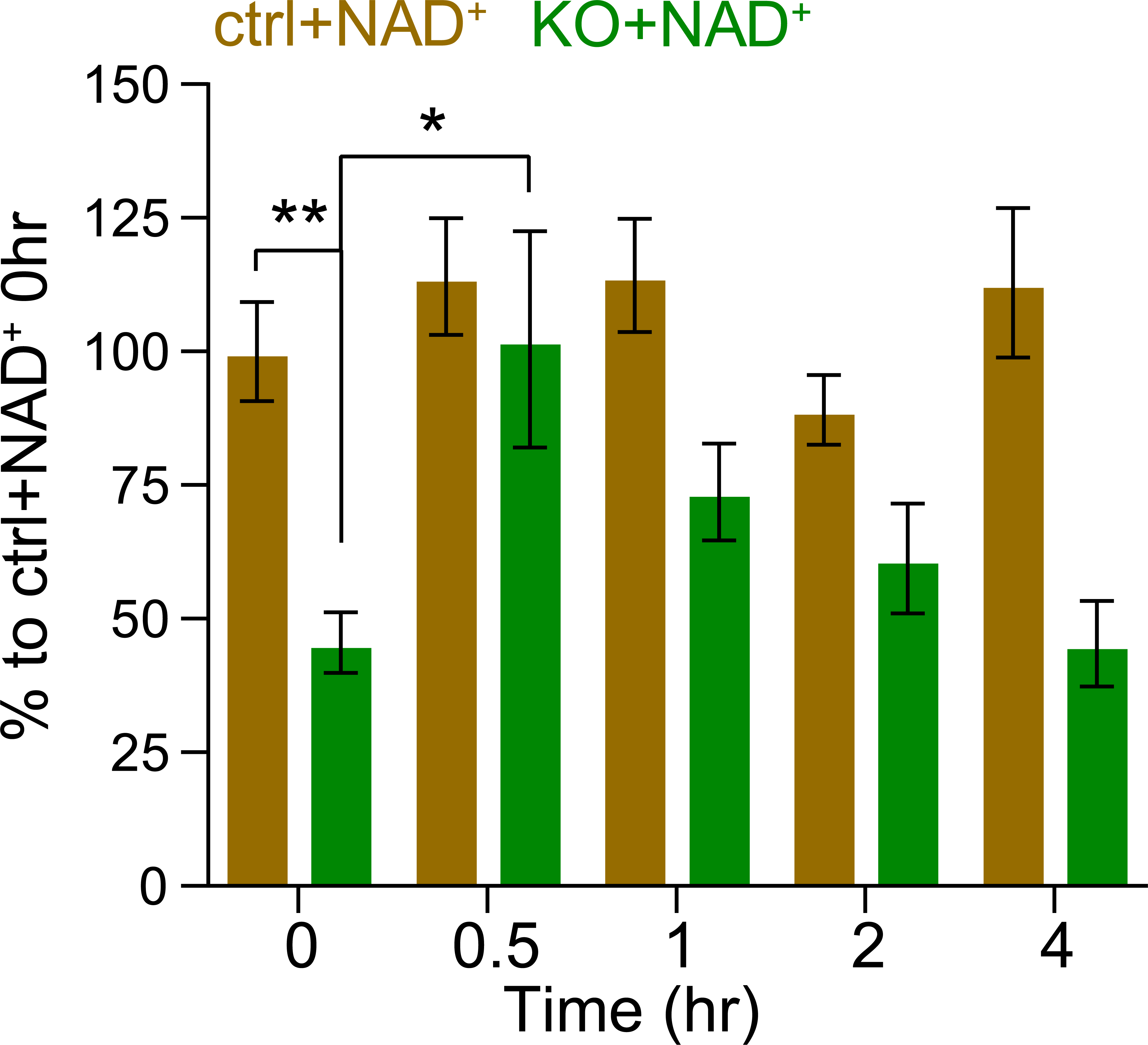
Increase of intracellular NAD by exogenous supplementation. The change of intracellular NAD^+^ abundance over time after one-time 1 mM NAD^+^ supplementation into the culture medium. All groups were normalized to ctrl+NAD^+^ 0 h time point and expressed as a percentage. Numbers (culture wells) and statistics: 13-14 ctrl and 12-14 KO from 4 independent experiments, Kruskal-Wallis test with Dunn’s multiple comparisons test was used, *p= 0.049, **p= 0.0023.

**Fig 6-S1.**
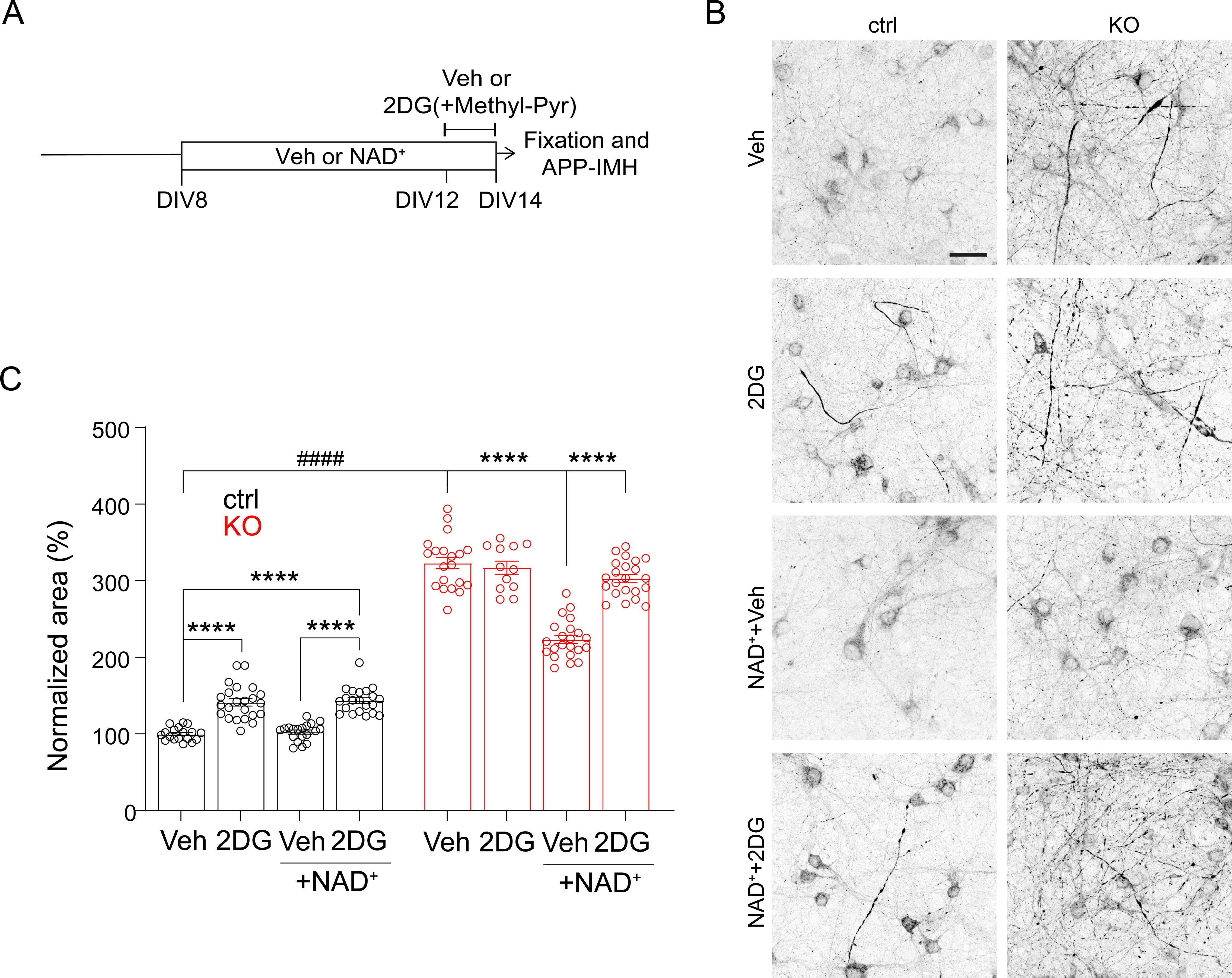
NAD supplementation depends on glycolysis to ameliorate APP accumulation in NMNAT2 KO neurons. (**A**) Timeline of NAD^+^ supplementation, 2DG(+Methyl-pyr) treatment, and immunostaining. (**B**) Representative confocal images of APP staining of DIV14 ctrl or KO neurons under Veh (vehicle) or NAD^+^ supplementation, treated with Veh or 2DG. (**C**) Area of accumulated APP in DIV14 ctrl or KO neurons supplemented with Veh or NAD^+^ and treated with Veh or 2DG. All values were normalized to ctrl supplemented and treated with Veh. Numbers (images) and statistics: 18 ctrl+Veh, 20 KO+Veh, 23 ctrl+ 2DG, 12 KO+2DG, 19 ctrl+NAD^+^, 22 KO+NAD^+^, 21 ctrl+NAD^+^+2DG, and 22 KO+NAD^+^+2DG, Two-way ANOVA with Tukey’s multiple comparisons test was used, ****p<0.0001, ^####^p<0.0001. Except for KO+2DG, all groups were from 2 independent experiments. All above data represent mean ± SEM.

**Fig 7-S1.**
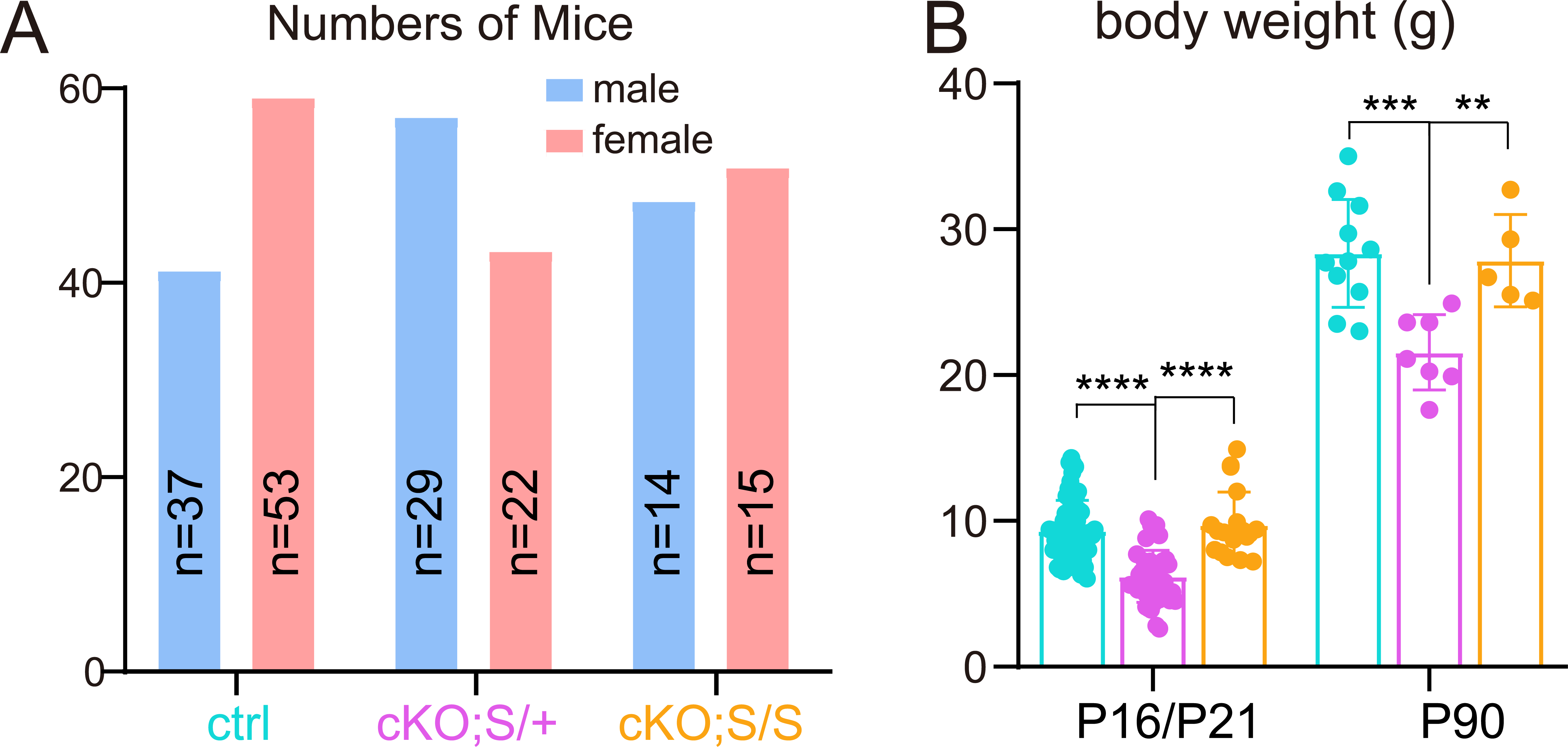
Gender ratio and body weight in surviving NMNAT2 cKO mice lacking one or two copies of SARM1. (**A**) Gender ratios in postnatal ctrl, NMNAT2 cKO; S^null^/+, and NMNAT2 cKO; S^null^/S^null^ mice pooled from progenies derived from various parent genotypes combinations. (**B**) Body weight of ctrl, NMNAT2 cKO; S^null^/+, and NMNAT2 cKO; S^null^/S^null^ mice at P16/21, and P90. Number of mice: P16/21, 61 ctrl, 34 NMNAT2 cKO; S^null^/+, 24 NMNAT2 cKO; S^null^/S^null^; P90, 11 ctrl, 7 NMNAT2 cKO; S^null^/+, 5 NMNAT2 cKO; S^null^/S^null^. Kruskal-Wallis test with two-stage linear step-up procedure of Benjamini, Krieger and Yekutieli was used for P16/21 data, ****p<0.0001. One-way ANOVA with two-stage linear step-up procedure of Benjamini, Krieger and Yekutieli was used for P90 data, (ctrl vs. NMNAT2 cKO; S^null^/+), ***, p= 0.0004, q=0.0004; (NMNAT2 cKO; S^null^/+ vs. NMNAT2 cKO; S^null^/S^null^), **, p= 0.0039, q= 0.002. All above data represent mean ± SEM.

**Fig 7-S2.**
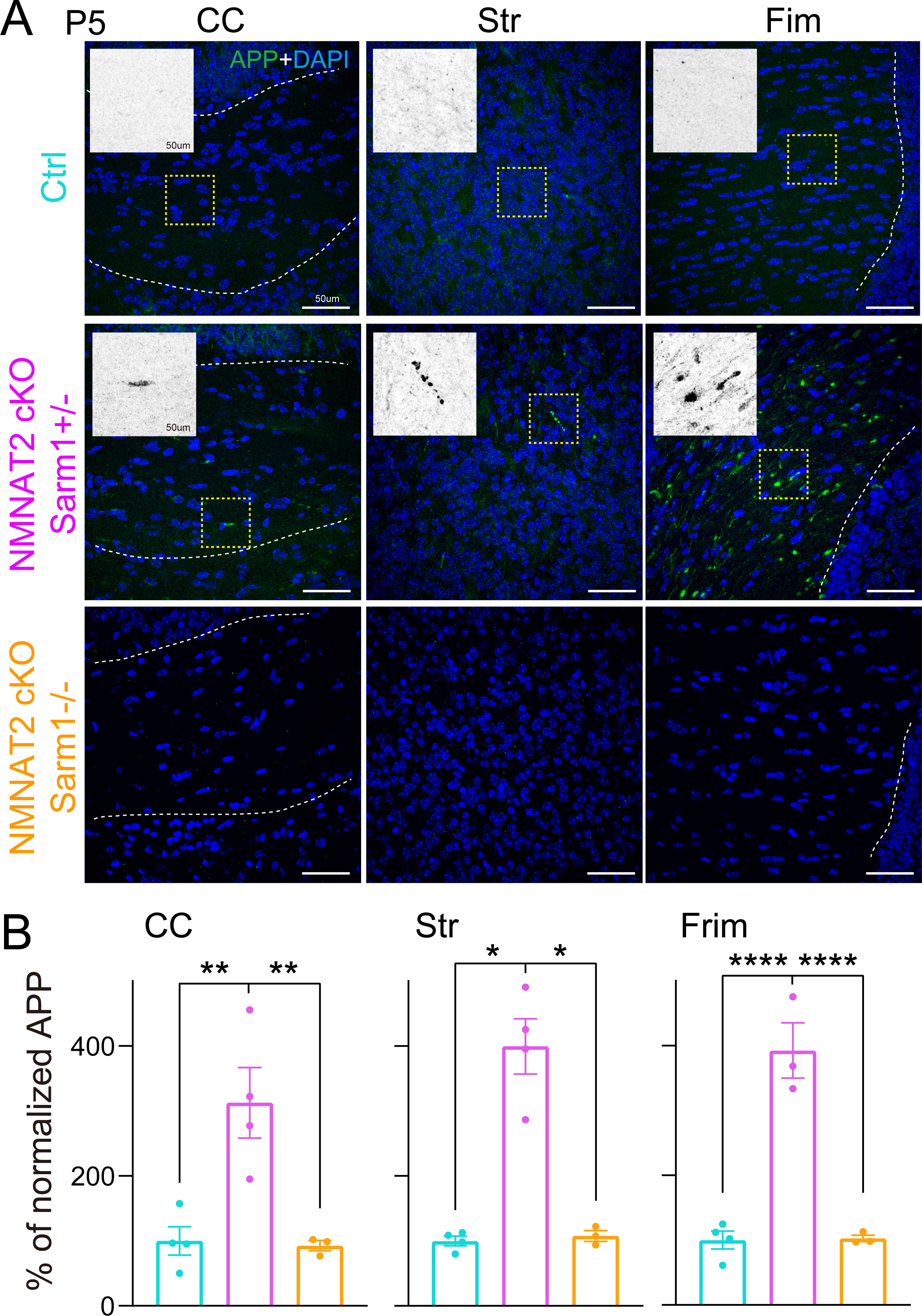
APP accumulation in P5 brains of NMNAT2 cKO mice lacking one or two copies of SARM1. (**A**) Immunohistology images showing APP accumulation in the corpus callosum, fimbria, and striatum of P5 NMNAT2 cKO; S^null^/+, but not in ctrl and NMNAT2 cKO; S^null^/S^null^ brains. (**B**) Normalized APP signals in P5 brains. Number of mice: 4 ctrl, 4 NMNAT2 cKO; S^null^/+, 3 NMNAT2 cKO; S^null^/S^null^. CC, corpus callosum; Str, striatum; Fim, Fimbria. For CC, one-way ANOVA with two-stage linear step-up procedure of Benjamini, Krieger and Yekutieli was used, (Ctrl vs. NMNAT2 cKO; S^null^/+) **, p=0.0032, q= 0.0021; (NMNAT2 cKO; S^null^/+ vs. NMNAT2 cKO; S^null^/S^null^) **, p=0.004, q=0.0021. For Str, Kruskal-Wallis test with two-stage linear step-up procedure of Benjamini, Krieger and Yekutieli was used, (Ctrl vs. NMNAT2 cKO; S^null^/+) *, p=0.0142, q=0.0298; (NMNAT2 cKO; S^null^/+ vs. NMNAT2 cKO; S^null^/S^null^) *, p=0.0414, q= 0.0435. For Frim, one-way ANOVA with two-stage linear step-up procedure of Benjamini, Krieger and Yekutieli was used, ****p<0.0001. All above data represent mean ± SEM.

**Fig 8-S1.**
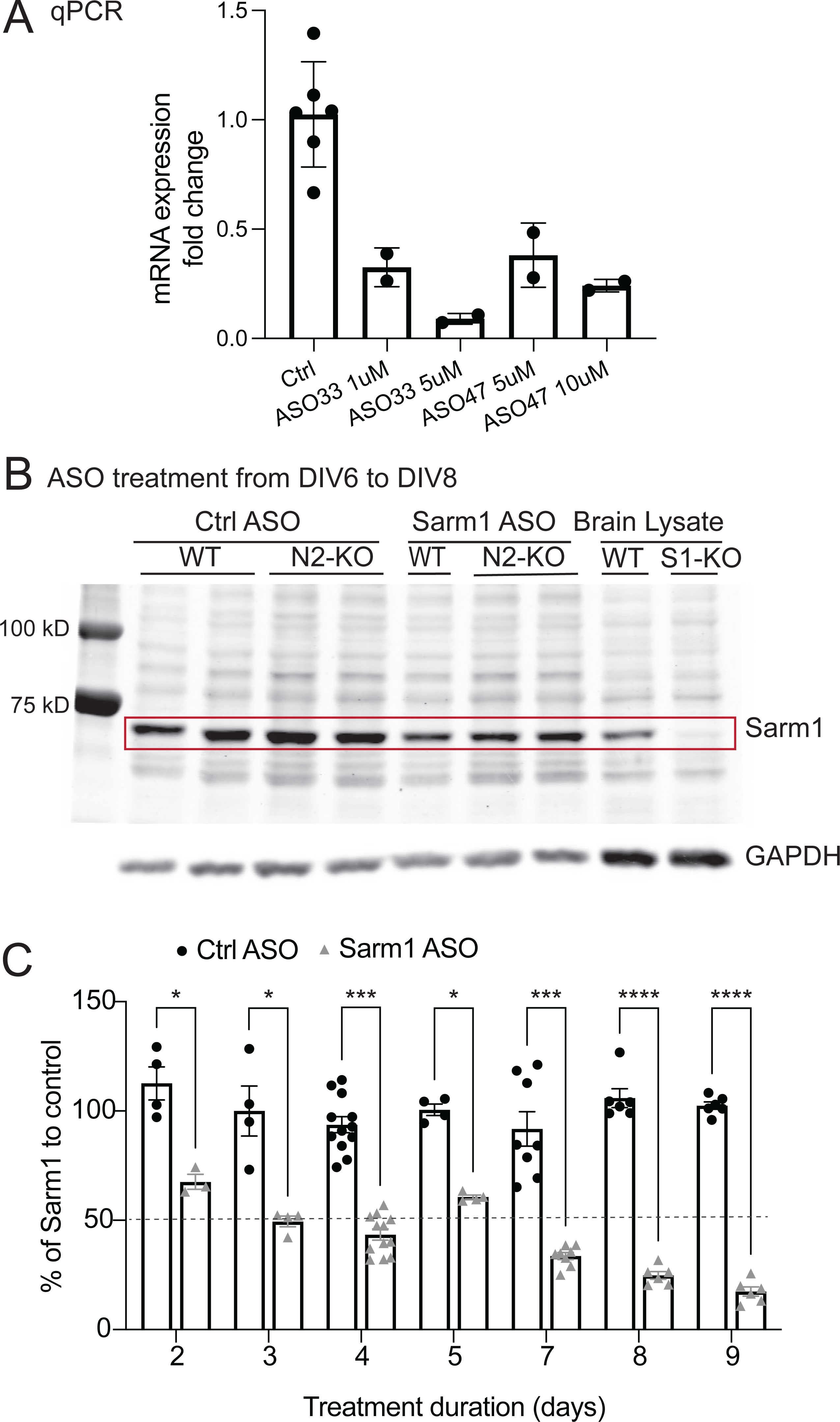
SARM1 knockdown validation. (**A**) RT-qPCR quantification of sarm1 mRNA levels across ASO treatments in primary mouse cortical neurons. Cells were treated for 2 days (DIV5-7) with a control or a SARM1 ASO (33 or 47) at two different concentrations to test efficacy. Number of wells: 6 ctrl; 2 ASO33 at 1 uM or 5 uM, 2 ASO47 at 5 uM or 10 uM. (**B**) Representative western blotting detecting SARM1 and GAPDH (internal control) protein levels at DIV8 after ASO treatment since DIV6. Brain lysates of SARM1 WT and SARM1-null (S^null^/S^null^) mice were included for antibody specificity validation. (**C**) Quantification of SARM1 protein level across various ASO treatment duration, as detected by western blot. SARM1 protein level was normalized to GAPDH first, then normalized to its corresponding ctrl-ASO-treated group. Numbers of wells: 2 days Sarm1 ASO, n=3. 2 days Ctrl ASO, 3 days Ctrl ASO, 3 days Sarm1 ASO, 5 days Ctrl ASO, 5 days Sarm1 ASO, n=4.8 days Ctrl ASO, 8 days Sarm1 ASO, 9 days Ctrl ASO, 9 days Sarm1 ASO, n=6. 7 days Ctrl ASO, 7 days Sarm1 ASO, n=8. 4 days Ctrl ASO, 4 days Sarm1 ASO, n=12. Statistics were conducted with the Kruskal-Wallis test and corrected for multiple comparisons by controlling the False Discovery Rate using the Two-stage step-up method of Benjamin, Krieger, and Yekutieli. All above data represent mean ± SEM. *, q<0.05, ***, q<0.001, ****, q<0.0001.

**Fig 8S-2.**
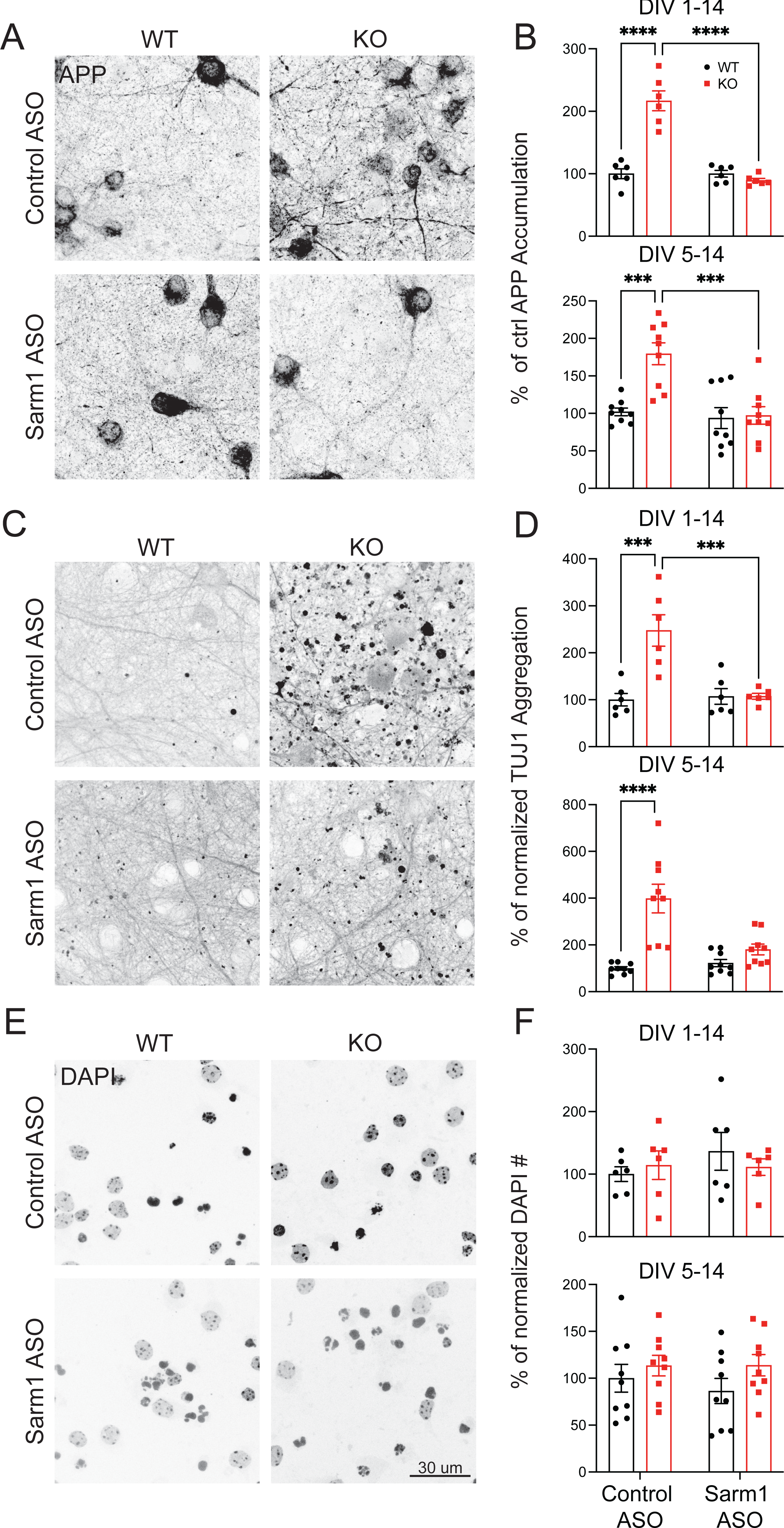
SARM1 knockdown prevents APP accumulation and axon degeneration in NMNAT2 KO neurons in vitro. (**A**) Representative images illustrating the effects of ctrl-ASO and SARM1-ASO on APP accumulation in WT and NMNAT2 KO primary cultured cortical neurons. (**B**) Quantification of APP accumulation area following ASO treatment of two timelines: DIV1 to DIV14 and DIV5 to DIV14. (**C**) Representative images show TUJ1 axonal staining of WT and KO cells following treatment with ctrl-ASO and SARM1-ASO. (**D**) Percentage of the total TUJ1 area found as aggregates following control and SARM1 ASO (ASO 33) treatments at DIV1-14 and DIV5-14. (**E**) Representative images of DAPI staining and quantification (**F**) of live cells in every sampled image. Image numbers: DIV1-14, N=6 for every group, from 2 independent experiments; DIV5-14, N=9 for every group, from 3 independent experiments. 3 images were collected from each experimental batch. Data for panel D, DIV5-14 was not normally distributed, so a Kruskal-Wallis test multiple comparisons test was performed. All other data sets were analyzed using two-way ANOVA and Tukey multiple comparisons. All above data represent mean ± SEM. ***, p<0.001,****, p<0.0001.

